# Exogenous Photoreceptor-Specific N-Glycosylated PROM1 Rescues Retinal Degeneration in Patient and Mouse Models

**DOI:** 10.1101/2025.08.06.668899

**Authors:** Ping Xu, Fuying Guo, Yuan Wang, Guifu Chen, Xiaojing Song, Bella Y. Luo, Dandan Zheng, Guanjie Gao, Wenjing Yin, Suai Zhang, Bruce T. Lahn, Xiufeng Zhong

**Affiliations:** State Key Laboratory of Ophthalmology, Zhongshan Ophthalmic Center, Sun Yat-sen University, Guangdong Provincial Key Laboratory of Ophthalmology and Visual Science, Guangzhou, 510060, China; Lantu Biopharma, Guangzhou, China; VectorBuilder, Chicago, IL, USA; Future Institute of Gene Delivery Research, Guangzhou, China

**Author notes:** Correspondence: Xiufeng Zhong, State Key Laboratory of Ophthalmology, Zhongshan Ophthalmic Center, Sun Yat-sen University, 54 S. Xianlie Road, Guangzhou 510060, China. These authors have contributed equally to this work (Ping Xu, Fuying Guo, Yuan Wang).

## Abstract

PROM1 is widely expressed across various tissues. However, its pathogenic mutations are exclusively associated with inherited retinal dystrophy (IRD). The mechanisms underlying this retina-specific vulnerability remain poorly understood, and no effective treatment currently exists for PROM1-IRD. Here, we utilized urine cells, hiPSCs, hiPSC-RPE cells, retinal organoids (ROs) and *Prom1^-/-^* mice to address these challenges. During photoreceptor development in ROs, PROM1 co-localized with ciliary marker ARL13B and outer segment (OS) marker PRPH2. It exhibited photoreceptor-specific mRNA splicing isoforms and unique N-glycosylation. In IRD patient-specific models with the PROM1 c.619G>T (p.E207X) homozygous mutation, we observed nonsense-mediated mRNA decay and altered splicing, leading to complete loss of PROM1 protein and OS-like structure disruption, faithfully recapitulating PROM1-IRD pathology. To rescue these defects, we engineered a photoreceptor-specific *AAV7m8-CRXp-hPROM1*, which successfully restored PROM1 expression and OS-like structures in patient-derived ROs. Therapeutic efficacy was further validated in *Prom1^-/-^* mice, where subretinal delivery of *AAV8-CRXp-hPROM1* led to photoreceptor-specific expression of human PROM1, significantly preserving OS morphology and improving visual function. These findings not only provide the first solid preclinical evidence supporting gene therapy for PROM1-IRD, but also reveal photoreceptor-specific vulnerability to PROM1 mutations, offering a novel conceptual framework for investigating and treating related IRDs.

## Introduction

Human prominin-1 (PROM1, also known as AC133 or CD133) is a pentaspan transmembrane glycoprotein broadly expressed on plasma membrane protrusions (1–3). It is widely recognized as a marker of hematopoietic and cancer stem cells (4, 5). PROM1 participates in various cellular processes, including membrane morphogenesis, cell proliferation, apoptosis, autophagy, and epithelial-mesenchymal transition (EMT) (3, 6–8). To date, approximately 150 pathogenic variants in *PROM1* have been documented in the Human Gene Mutation Database (9). Paradoxically, despite its broad tissue distribution, mutations in *PROM1* are exclusively associated with a single-organ disorder known as PROM1-related inherited retinal dystrophy (PROM1-IRD, OMIM 604365), which can be inherited in either an autosomal dominant or recessive manner and manifests as cone-rod dystrophy, retinitis pigmentosa, or Stargardt-like disease (10, 11).

In retinal photoreceptors, PROM1 localizes to the outer segment (OS) and plays a critical role in disc morphogenesis (9, 12, 13). This photoreceptor-specific function has been partly attributed to alternative splicing of *PROM1* transcripts. The human *PROM1* gene harbors six alternative promoters (14), generating photoreceptor-specific isoforms mainly characterized by the inclusion of exon 4 and the exclusion of exons 26 and/or 25(9). In addition, PROM1 contains nine N-glycosylation sites located within its two large extracellular loops, each exceeding 250 amino acids (15, 16). Glycosylation is known to modulate protein function; for example, high mannose PROM1 facilitates interaction with DNMT1 to maintain the slow-cycling state and tumorigenic potential of glioma stem cells (17). However, whether PROM1 glycosylation contributes to its specialized role in photoreceptors remains to be elucidated.

In recent years, gene augmentation therapy has emerged as a promising therapeutic strategy for IRDs, offering the potential to restore the function of defective genes and achieve durable therapeutic outcomes. Among various gene delivery systems, adeno-associated virus (*AAVs)* have gained widespread application due to their genetic tractability, favorable safety profile, low immunogenicity, and ability to mediate sustained transgene expression following a single administration (18). A notable milestone in this field is Luxturna, the first FDA-approved *AAV2*-mediated gene therapy for RPE65-IRD (19). Encouraged by this success, gene augmentation therapies targeting other IRDs pathogenic genes such as *CYP4V2*, *ABCA4*, *BEST1*, and *CEP290* are currently under active preclinical or clinical investigation (20–23). Despite *PROM1* being a relatively prevalent IRDs pathogenic gene, therapeutic development for PROM1-IRD remains limited (24).

*Prom1* knockout (*Prom1^-/-^*) mice have been established and recapitulate key features of human PROM1-IRD, including progressive visual decline and retinal degeneration (25–29). Ultrastructural analyses using transmission electron microscopy have further revealed disorganized and malformed disc membranes in the photoreceptor outer segments of these models (25–28). In parallel, retinal organoids (ROs) derived from pluripotent stem cells recapitulate photoreceptor outer segments and exhibit light-responsive electrophysiological properties *in vitro* (30, 31). Together, these complementary model systems provide robust and translationally relevant platforms for the in-depth investigation of disease mechanisms and the preclinical evaluation of gene augmentation strategies for PROM1-IRD.

In this study, we investigated the role of PROM1 in photoreceptor development and the pathogenesis of PROM1-IRD, and evaluated the feasibility of gene augmentation therapy. Using urine cells (UCs), hiPSCs, hiPSC-RPE cells and ROs, we systematically characterized PROM1 expression, subcellular localization, and photoreceptor-specific mRNA alternative splicing and protein glycosylation. In parallel, patient-specific hiPSCs and ROs carrying a *PROM1* c.619G>T (p.E207X) homozygous mutation exhibited significant nonsense-mediated mRNA decay, aberrant splicing, and defects in OS-like structures. This structural impairment was recapitulated in a newly generated *Prom1^-/-^* mouse. To rescue PROM1-IRD, we developed a gene augmentation strategy using a modified *CRX* promoter to drive human PROM1 expression. This approach successfully restored OS morphology in both patient ROs and *Prom1^-/-^* mice. Collectively, our findings not only provide the first preclinical validation for gene therapy targeting PROM1-IRD, but also reveal photoreceptor-specific vulnerability to PROM1 mutations, offering a novel conceptual framework for investigating and treating related IRDs.

## Results

### Spatiotemporal expression pattern of PROM1 during photoreceptor development in ROs

Although the expression of PROM1 in the adult human retina is established, its spatiotemporal pattern during retinal development remains poorly understood (32). Here, we utilized ROs, which recapitulate key stages of human retinal development *in vitro* (30, 31), to systematically characterize the dynamic expression of PROM1. At the early stage (day 24, D24) of wild-type ROs (C1-ROs), immunofluorescence revealed a punctate distribution of PROM1 broadly expressed in both VSX2⁺ retinal progenitor cells (RPCs) and VSX2⁻ cells (Fig. S1A), suggesting non-specific expression at this stage. By D40 and D60, PROM1 became enriched at the apical membrane of RPCs and photoreceptor precursors, showing clear colocalization with the ciliary marker ARL13B (Fig. 1A). Although a subset of cells began expressing the pan-photoreceptor marker Recoverin (RCVRN) at D60, PROM1 expression remained comparable between RCVRN^+^ cells and adjacent precursors (Fig. 1A; Fig. S1B).

**Figure 1.**
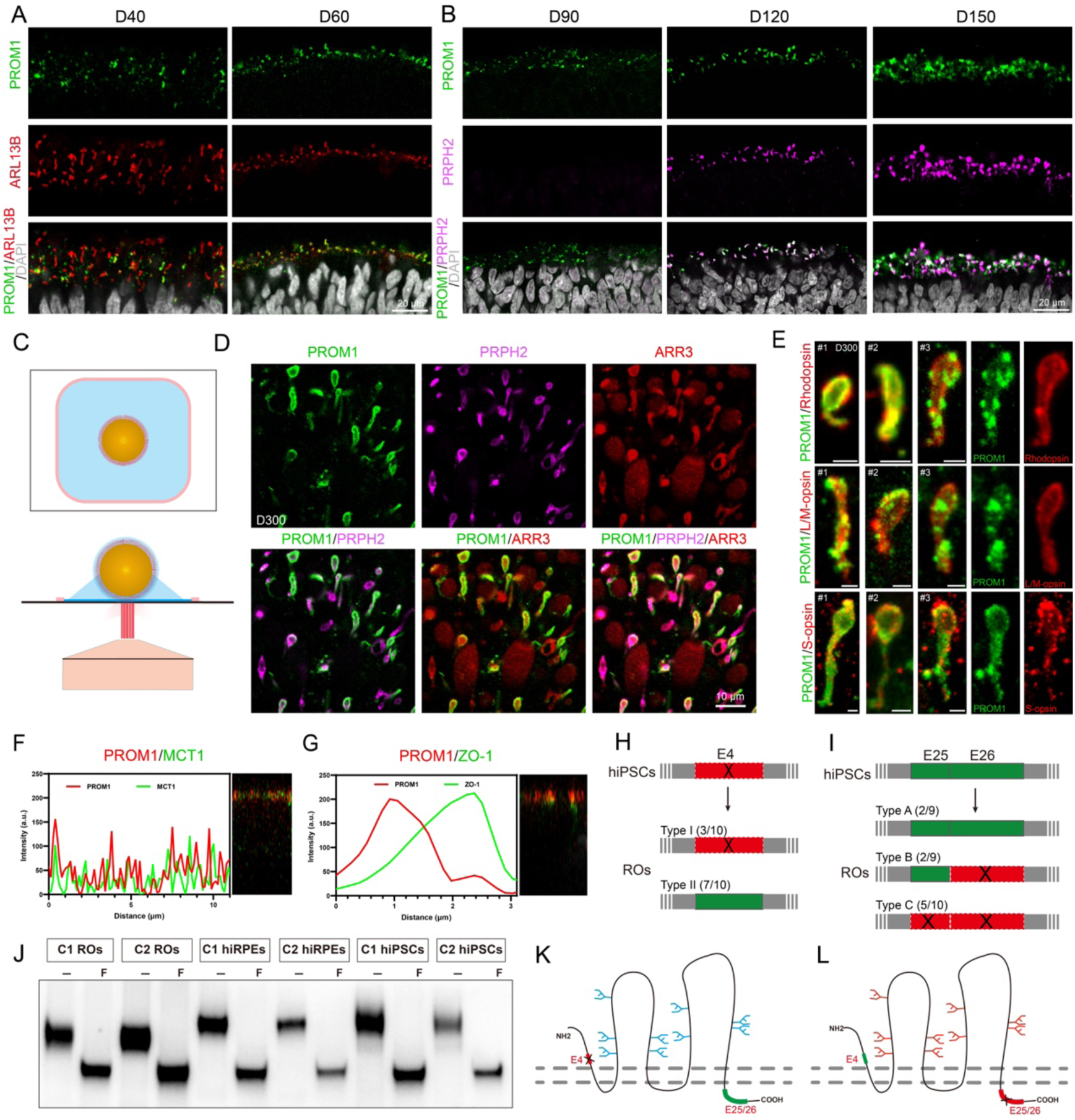
Photoreceptors Express a Distinct N-Glycosylated PROM1 in ROs. **(A and B)** Immunostaining of PROM1 during C1-ROs differentiation, co-labeled with ARL13B (cilia marker) or PRPH2 (OS marker). Nuclei are counterstained with DAPI. Time points: D40, D60, D90, D120, and D150. Scale bar, 20 µm. **(C)** Schematic diagram of whole-ROs imaging. The top view illustrates the coverslip (black square), the hydrophobic barrier drawn with a PAP pen (pink square), the antifade mounting medium (blue area), and the RO (yellow sphere). The frontal view shows the red laser and the confocal microscope objective positioned beneath the coverslip. **(D)** Whole-mount staining of D300 C1-ROs showing PROM1 expression in PRPH2^+^ OS-like structures and ARR3^+^ cones. Scale bar, 10 µm. **(E)** Co-localization of PROM1 with Rhodopsin (rods), L/M-opsin (L/M-cones), and S-opsin (S-cones) in D300 C1-ROs. Scale bar, 2 μm. **(F and G)** Confocal immunostaining of PROM1, MCT1, and ZO-1 in C1 hiPSC-RPE cells cultured on Transwells. Y–Z projections illustrate subcellular localization. **(H and I)** Schematics showing alternative splicing patterns of *PROM1* in exon 4 (H) and exons 25–26 (I) in C1-hiPSCs versus C1-ROs. **(J)** Western blot analysis of PROM1 expression and N-glycosylation in hiPSCs, hiPSC-RPE cells, and ROs. C1 and C2 represent different normal individuals. “F” indicates Glycopeptidase F-treated samples; “−” denotes untreated controls. **(K and L)** Diagrams depicting glycosylated PROM1 in the plasma membrane of hiPSCs (K) and photoreceptors (L).

At D90, PROM1 expression persisted in photoreceptors, despite the absence of PRPH2 expression (Fig. 1B; Fig. S1B). By D120, cones marker Arrestin3 (ARR3) and rods marker Rhodopsin (RHOD) were detected and displayed an organized distribution in the outer layer of the C1-ROs (Fig. S1C). At this stage, PRPH2 expression also emerged and colocalized with PROM1 in OS-like structures (Fig. 1B). This colocalization pattern remained consistent from D150 to D270 (Fig. 1B; Fig. S2A). The observed spatiotemporal expression profile was further confirmed in an independent wild-type line (C2-ROs) at D60, D90, D120, and D150 (Fig. S2B).

Given the distinct morphologies and proteins requirements for rods and cones OS formation (33), we next examined whether PROM1 was expressed across all photoreceptor subtypes. Cryosections of C1-ROs from D120 to D210 were analyzed (Fig. 1B; Fig. S2A). However, the structural integrity of OS-like structures was significantly compromised due to dehydration, freezing, sectioning and rehydration during immunostaining of cryosections, limiting the resolution of PROM1 subcellular localization. To address this issue, we optimized a whole-mount immunostaining protocol for intact ROs, which better preserves the OS-like structures. Specifically, ROs were subjected only to paraformaldehyde fixation and serum blocking prior to staining, therapy minimizing damage from extensive experimental procedures, particularly freezing and rehydration (Fig. 1C). Confocal imaging of D300 C1-ROs revealed robust colocalization of PROM1 with both PRPH2 and ARR3 within intact OS-like structures (Fig. 1D). Moreover, PROM1 expression was detected in all major photoreceptor subtypes, including rods (RHOD⁺), L/M-cones (LM-opsin⁺), and S-cones (S-opsin⁺), supporting its broad and subtype-independent expression in human photoreceptors (Fig. 1E).

To further assess PROM1 expression in retinal pigment epithelial (RPE) cells, C1 hiPSC-RPE cells were isolated from D45 C1-ROs and cultured on Transwell inserts for 6 weeks. Phase-contrast microscopy showed a typical cobblestone-like morphology with abundant pigmentation (Fig. S3A). Immunofluorescence analysis demonstrated that PROM1 localized above the tight junction protein ZO-1 and colocalizing with the apical membrane marker MCT1 (Fig. 1F-G; Fig. S3B), indicating its enrichment in the apical microvilli of C1 hiPSC-RPE cells.

### *PROM1* exhibits specific mRNA alternative splicing isoforms and protein N-glycosylation patterns in photoreceptors

Although PROM1 is broadly expressed across various tissues, pathogenic variants selectively affect photoreceptors. To determine whether photoreceptors express distinct *PROM1* isoforms compared to other cell types, *PROM1* mRNA variants were analyzed in hiPSCs and ROs. Notably, the D270 C1-ROs used for analysis preserved only the translucent outer region enriched in photoreceptors. To ensure consistent annotation, exon numbering was based on *PROM1* transcript variant 1 (NM_006017.3), the longest isoform comprising all 28 exons (Fig. S4A). RT-PCR was performed to amplify regions spanning the coding sequences of exons 3–12 and 24–27 (Fig. S4B). Sanger sequencing showed that C1-hiPSCs predominantly exhibited exon 4 skipping and inclusion of exons 25–26, whereas C1-ROs displayed overlapping peaks in these regions, indicating mixed isoform populations (Fig. 1H–I; Fig. S4C). TA cloning and sequencing of RT-PCR products from C1-ROs confirmed that most clones retained exon 4 (7/10), while a minority showed exon 4 skipping (3/10). For exons 25–26, two clones retained both exons (2/9), two retained exon 25 but skipped exon 26 (2/9), and the majority lacked both exons (5/9) (Fig. 1I; Fig. S4D–E). These photoreceptor-specific alternative splicing isoforms may underlie the unique role of PROM1 in OS formation and maturation.

In addition to transcriptional diversity at the mRNA level, glycosylation, a post-translational modification, can significantly influence protein folding, localization and function (7). To determine whether PROM1 exhibits tissue-specific glycosylation in photoreceptors, we compared its glycosylation patterns in hiPSCs, hiPSC-RPE cells, and ROs. Western blot analysis confirmed the presence of PROM1 across all three cell types. Notably, PROM1 from D3 hiPSCs and 6-week hiPSC-RPE cells displayed a higher apparent molecular weight compared to that from D150 and D270 ROs (Fig. 1J; Fig. S5; Fig. S11D). Following treatment with PNGase F, an enzyme that removes N-linked glycans, the molecular weight discrepancies were markedly diminished, indicating that PROM1 in photoreceptors within ROs undergoes distinct, tissue-specific N-glycosylation (Fig. 1J; Fig. S5).

### The *PROM1* c.619G>T mutation causes autosomal recessive IRD

To explore the genotype-phenotype relationship in PROM1-IRD, a family affected by early-onset retinal dystrophy was enrolled in this study (Fig. 2A). The proband (F2-2) developed progressive bilateral visual impairment and photophobia beginning at age 9. At his initial ophthalmic examination at age 13, best-corrected visual acuity (BCVA) was 0.4 in both eyes. Fundus examination revealed well-demarcated, oval-shaped brown lesions (∼0.7 disc diameters) in the macula surrounded by a depigmented halo. Fundus fluorescein angiography (FFA) showed a classic bull’s-eye pattern in both eyes (Fig. S6A). By age 35, BCVA had declined to 0.04/0.06, and fundus color photography demonstrated a triangular-shaped central macular lesion with perifoveal pigment clumping (Fig. S6C). At age 38, visual acuity had further deteriorated to hand motion in both eyes. Fundus examination revealed stable macular pathology, accompanied by bone spicule-like pigmentation and chorioretinal atrophy in peripheral retina (Fig. 2B; Fig. S6D-E). FFA and indocyanine green angiography confirmed the fundus lesions and did not observe any neovascular leakage (Fig. 2C, Fig. S6F–I). Blue autofluorescence imaging showed markedly reduced signal intensity in the posterior pole (Fig. S6B). Spectral-domain optical coherence tomography (SD-OCT) demonstrated outer retinal atrophy and cystoid changes in the macular region (Fig. 2D). Flash electroretinography (fERG) revealed extinguished dark-adapted and light-adapted responses in both eyes (data not shown).

**Figure 2.**
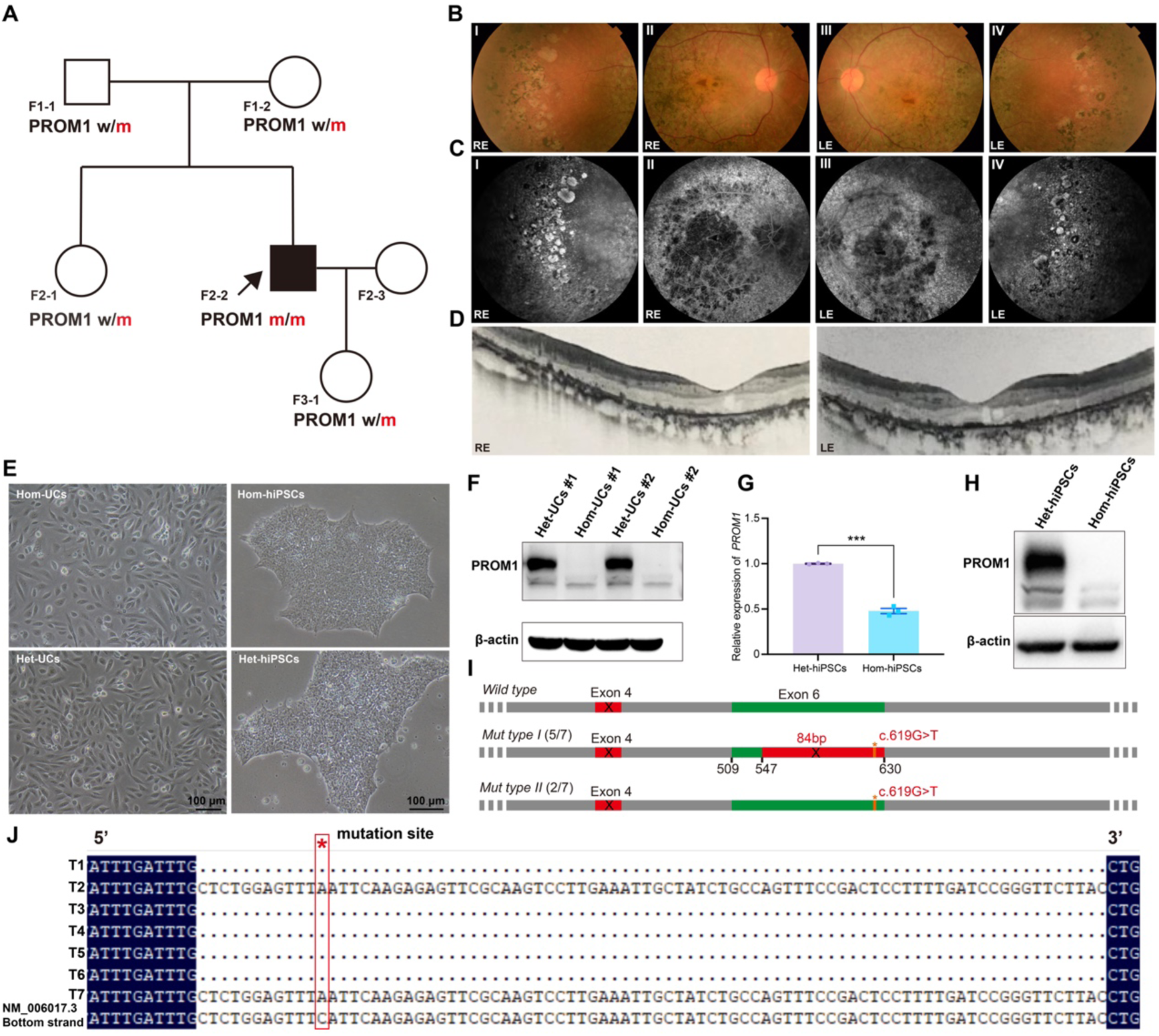
*PROM1* c.619 G>T mutation induces NMD and NAS. **(A)** Pedigree of the PROM1-IRD family. The proband (F2-2) indicated by a black arrow. **(B-D)** Multimodal retinal imaging of the 35-year-old proband, including fundus photographs(B), FFA (C) and SD-OCT (D). RE, right eye; LE, left eye. **(E)** Bright-field images of UCs and hiPSCs from the proband (Hom) and his mother (Het). Scale bar, 100 μm. **(F and H)** Western blot showing PROM1 protein levels in UCs and hiPSCs. β-actin serves as the loading control. **(G)** qRT-PCR analysis of *PROM1* mRNA expression in Het-hiPSCs (n=3) and Hom-hiPSCs (n=3). *Error bars represent mean ± S.E.M.; ***P < 0.001.* **(I and J)** TA cloning and sequencing of *PROM1* cDNA from Hom-hiPSCs. Among 7 clones, 5 exhibited an 82 bp deletion (Mut type I), and 2 harbored the c.619 G>T point mutation (Mut type II). The corresponding sequence is detailed in (J).

Next-generation sequencing using a retinal disease gene panel covering 381 known causative genes identified a *PROM1* c.619G>T (p.E207X) homozygous mutation (Fig. S7A). Based on the clinical phenotype and the established link between *PROM1* and IRDs (34, 35), a diagnosis of PROM1-IRD was confirmed. Sanger sequencing of family members confirmed heterozygous carrier status in the proband’s parents, sister, and daughter (Fig. 2A; Fig. S7A). All carriers exhibited normal BCVA and no retinal abnormalities upon examination, consistent with autosomal recessive inheritance of the mutation.

### The *PROM1* c.619G>T (p.E207X) mutation induces NMD and NAS mechanisms

Nonsense mutations constitute a substantial fraction of pathogenic variants in IRDs and are known to induce not only premature protein truncation but also trigger nonsense-mediated mRNA decay (NMD) and nonsense-associated altered splicing (NAS) (36, 37). In our previous study, UCs from a patient harboring a *PROM1* c.619G>T homozygous mutation and his heterozygous mother were reprogrammed into hiPSCs (38), hereafter referred to as Hom-hiPSCs and Het-hiPSCs, respectively (Fig. 2E). These hiPSC lines were used to investigate the effects of the mutation on *PROM1* mRNA and protein expression.

Bidirectional Sanger sequencing of *PROM1* cDNA from both UCs and hiPSCs revealed a prominent overlapping peak corresponding to an 84 bp region (c.547–630) encompassing the mutation site (c.619), with lower signal intensity in Het compared to Hom samples (Fig. 2H; Fig. S7B-C). To clarify the sequence composition of this region, PCR amplification of the corresponding cDNA segment from Hom-hiPSCs was followed by TA cloning. Among the seven clones analyzed, two (T2 and T7) carried the expected c.619G>T mutation, while the remaining five exhibited a consistent 84 bp deletion within exon 6, indicating mutation-induced aberrant splicing. In addition, qRT-PCR revealed significantly reduced *PROM1* mRNA in both Het-hiPSCs and Hom-hiPSCs compared to control C1-hiPSCs (Fig. 2G), suggesting the activation of NMD pathway. Consistently, Western blot using both N-terminal (AA 20–108) and C-terminal PROM1-specific antibodies failed to detect PROM1 protein in Hom-UCs and Hom-hiPSCs (Fig. 2F, H; Fig. S8), confirming a complete loss of protein expression.

### The *PROM1* mutation does not impair retinal development in Hom-ROs

ROs were generated from C1-hiPSCs, Het-hiPSCs, and Hom-hiPSCs lines using established protocols (30, 39). The patient-derived Hom-ROs exhibited proper laminar organization and contained all major retinal cell types, including ganglion cells (RBPMS⁺), photoreceptors (OTX2⁺ and RCVRN⁺), horizontal cells (PROX1⁺), amacrine cells (AP2α⁺), bipolar cells (PKCα⁺), and Müller glia (CRALBP⁺) (Fig. S9A). Additionally, both rods (Rhodopsin⁺) and cones subtypes labeled with L/M-opsin or S-opsin were properly differentiated and exhibited normal spatial organization within the outer layer of the ROs (Fig. S9B-C). These findings demonstrate that the *PROM1* c.619G>T homozygous mutation does not disrupt retinal cell type differentiation or spatial patterning.

Quantitative analysis of the outer nuclear layer (ONL) at day 150 revealed no appreciable differences in thickness among Hom-ROs, C1-ROs, and Het-ROs (Fig. 3K). Consistently, TUNEL staining showed similar proportions of photoreceptors apoptosis across the three groups (Fig. 3L-M), suggesting that the mutation does not elicit increased photoreceptors death under basal conditions. Intriguingly, bulk RNA-seq analysis revealed that the apoptosis-related PI3K-Akt signaling pathway (ko04151) was among the most significantly enriched pathways in Hom-ROs compared with both C1-ROs and Het-ROs (Fig. 3F; Fig. S10A, D). No such enrichment was observed between C1-ROs and Het-ROs (Fig. S10E), pointing to a homozygous mutation-specific transcriptional alteration. Notably, unlike observations in other tissues (40), most differentially expressed genes (DEGs) associated in this pathway were upregulated in Hom-ROs. Specifically, 55 DEGs were identified between Hom-ROs and C1-ROs, and 34 between Hom-ROs and Het-ROs, with 24 overlapping genes (21 upregulated, 3 downregulated) (Fig. 3G; Fig. S10B-C). Importantly, comparative analysis of the apoptosis pathway (ko04210) revealed no significant alterations among the three groups, further supporting the conclusion that the *PROM1* mutation does not induce photoreceptor apoptosis in Hom-ROs.

**Figure 3.**
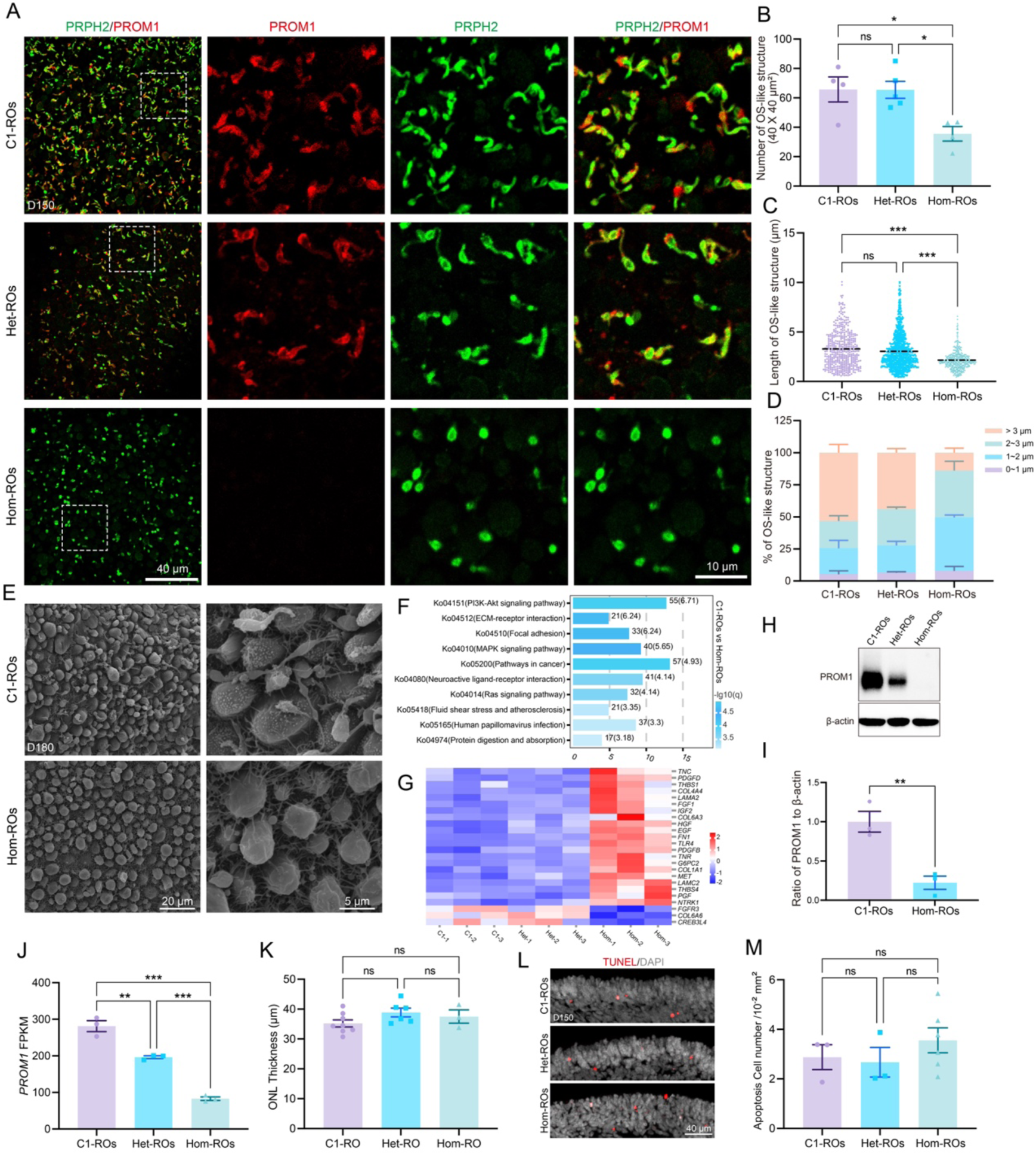
*PROM1* mutation impairs photoreceptors OS-like structures in ROs. **(A)** Immunostaining of PROM1 and PRPH2 in D150 ROs. Regions within white dashed boxes are shown enlarged. Scale bars: 40 μm (overview) and 10 μm (magnified). **(B and C)** Quantitative analysis of OS-like structures lengths in C1-ROs (n=444), Het-ROs (n=873), and Hom-ROs (n=396). The lengths were categorized into four groups: 0–1 μm, 1–2 μm, 2–3 μm, and >3 μm. **(D)** Quantification of OS-like structures per 40×40 μm². C1-ROs (n=4), Het-ROs (n=5), Hom-ROs (n=4). **(E)** TEM images of OS-like structures in D180 C1-ROs and Hom-ROs. Scale bars: 20 μm (left), 5 μm (right). **(F)** Top 10 significantly enriched KEGG pathways comparing D150 C1-ROs and Hom-ROs. **(G)** Heatmap of DEGs in the PI3K-Akt signaling pathway across all groups. **(H and I)** Western blot showing PROM1 protein levels in D150 ROs (n=4). β-actin as control. **(J)** RNA-seq analysis showing FPKM values of *PROM1* in D150 ROs (n=3). **(K)** ONL thickness quantification in D150 ROs based on DAPI staining. C1-ROs (n=8), Het-ROs (n=6), Hom-ROs (n=3). **(L and M)** TUNEL staining for photoreceptors apoptosis in D150 ROs. Apoptotic cells per 0.01 mm² quantified. C1-ROs (n=3), Het-ROs (n=3), and Hom-ROs (n=6); Scale bars, 40 μm. *Error bars represent mean ± S.E.M.; ns, not significant (P ≥ 0.05); *P < 0.05; **P < 0.01; ***P < 0.001*.

### *PROM1* mutation impairs OS-like structures in patient-specific Hom-ROs

PROM1 expression and localization were first assessed across the three groups of ROs. Bulk RNA-seq analysis showed significantly reduced *PROM1* FPKM values in both Het-ROs and Hom-ROs compared to C1-ROs, consistent with the finding in hiPSCs (Fig. 2G and Fig. 3J). Immunostaining of ROs from D40 to D270 demonstrated that Het-ROs exhibited PROM1 expression patterns similar to those in C1-ROs and C2-ROs, whereas PROM1 expression was undetectable in Hom-ROs (Fig. S11A-B). This absence was confirmed using two independent antibodies targeting the N- and C-terminal regions of PROM1. Both antibodies strongly labeled OS-like structures in Het-ROs but failed to detect PROM1 in Hom-ROs (Fig. S11C). Western blot further corroborated the complete loss of PROM1 in Hom-ROs (Fig. 3H-I; Fig. S11D).

To assess the effect of PROM1 mutation on photoreceptor outer segments, whole-mount staining was performed on D150 ROs. Confocal microscopy revealed that PRPH2-labeled OS-like structures in Hom-ROs lacked detectable PROM1 expression, in contrast to those in C1-ROs and Het-ROs (Fig. 3A). Quantitative analysis showed that both the length and number of OS-like structures were significantly reduced in Hom-ROs (Fig. 3B-C). Further stratification by length revealed a significant shift in the distribution of OS-like structures in Hom-ROs, with fewer structures exceeding 3 μm and a greater proportion falling within the 1–2 μm range (Fig. 3D). Scanning electron microscopy (SEM) further corroborated these findings (Fig. 3E). In addition, multiple punctate protrusions were observed on the inner segment membranes of C1-ROs, whereas the corresponding regions in Hom-ROs appeared smooth (Fig. 3E).

To further investigate the transcriptional alterations associated with these structural changes, gene ontology (GO) enrichment analysis was performed. Hom-ROs showed significant dysregulation of genes related to the plasma membrane (GO:0005886) and membrane microdomains (GO:0098857) compared to both C1-ROs and Het-ROs (Fig. S10F–H), supporting the impact of the PROM1 mutation on photoreceptor membrane organization and OS morphogenesis.

### PROM1 gene augmentation rescues photoreceptor OS-like structures in Hom-ROs

*AAV*-mediated retinal gene therapy is one of the most promising therapeutic strategies for autosomal recessive IRDs. To evaluate its feasibility for PROM1-IRD, we first optimized *AAV* transduction parameters in ROs. D60 C1-ROs, representing an early stage of photoreceptor development, were selected for initial testing. Four days after AAV infection, C1-ROs treated with *AAV7m8-CMV-GFP* exhibited stronger GFP fluorescence than those treated with *AAV2-CMV-GFP* or *AAV8-CMV-GFP*. However, they also displayed a loss of translucent morphology accompanied by extensive cell death, which was not observed in the untreated controls, indicating that D60 ROs are poorly tolerant to *AAV* infection (Fig. S12A-B). In contrast, D90 C1-ROs maintained normal morphology across all three AAV-treated groups. Among them, *AAV7m8-CMV-GFP* infected ROs showed significantly stronger GFP signals, supporting its superior transduction efficiency (Fig. 4A). Based on these findings, *AAV7m8* serotype was selected for subsequent experiments, and ROs older than 90 days were prioritized to minimize *AAV*-induced cytotoxicity.

**Figure 4.**
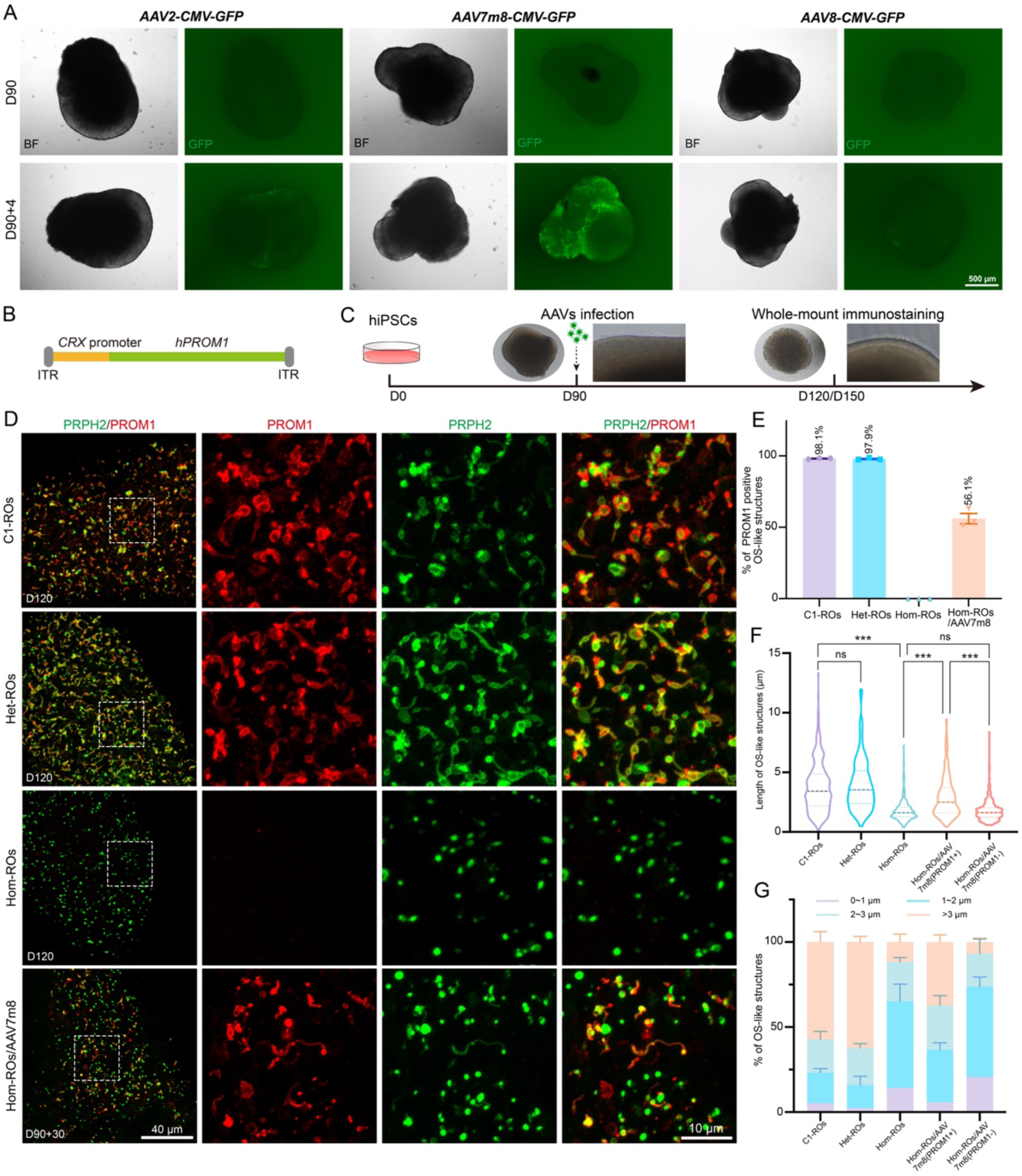
Gene augmentation rescues patient-specific photoreceptors phenotypes in ROs. **(A)** D90 C1-ROs was infected with *AAV2-CMV-GFP*, *AAV7m8-CMV-GFP or AAV8-CMV-GFP* (1×10^11^ vg/RO). Bright-field and fluorescence images taken before and after infection. Scale bar, 500 µm. **(B)** Schematic of *pAAV-CRXp-hPROM1* construct. **(C)** Experimental outline of the *AAV*-mediated gene augmentation in Hom-ROs. **(D)** Immunostaining for PROM1 and PRPH2 in D120 ROs. Hom-ROs infected at D90 is denoted Hom-ROs/AAV7m8. Regions within dashed boxes were magnified on the right. Scale bars: 40 µm (left) and 10 µm (right). **(E)** Proportions of PROM1^+^/PRPH2^+^ versus PROM1^-^/PRPH2^+^ OS-like structures in D90+30 Hom-ROs/AAV7m8. **(F and G)** Quantitative analysis of OS-like structures lengths in C1-ROs (n=486), Het-ROs (n=358), Hom-ROs (n=433), Hom-ROs/AAV7m8 (PROM1^+^/PRPH2^+^; n=297), and Hom-ROs/AAV7m8 (PROM1^-^/PRPH2^+^; n=242). *Error bars represent mean ± S.E.M.; ns, not significant (P ≥ 0.05); ***P < 0.001*.

To selectively expresses exogenous PROM1 in photoreceptors, a recombinant plasmid (*pAAV-CRXp-hPROM1*) was constructed using a modified *CRX* promoter (Fig. 4B). D90 Hom-ROs was subsequently infected with *AAV7m8-CRXp-hPROM1* at a dose of 1×10¹¹ vg per RO (referred to as the Hom-ROs/AAV7m8 group). One month post-infection, exogenous PROM1 was detected in a subset of PRPH2⁺ OS-like structures, with a transduction efficiency of approximately 56% (Fig. 4C-E). Quantitative analysis revealed that OS-like structures in the Hom-ROs/AAV7m8 were significantly longer than those in untreated Hom-ROs. Moreover, within the treated group, PROM1⁺ structures were notably longer than PROM1⁻ ones, and the proportion of structures exceeding 3 μm was significantly higher among the PROM1⁺ population (Fig. 4F-G, Fig. S13A).

Following an additional month of ROs culture, immunostaining confirmed that exogenous PROM1 expression persisted and remained localized to photoreceptor OS-like structures (Fig. S13B). Similarly, in D150 Hom-ROs examined 80 days after *AAV7m8-CRXp-hPROM1* infection, PROM1 expression was still detectable (Fig. S13C). These findings demonstrate sustained transgene expression and underscore the therapeutic potential of *AAV*-mediated gene augmentation for PROM1-IRD.

### Exogenous expression of human PROM1 ameliorates structural and functional impairments in *Prom1^-/-^* retinas

To further evaluate the feasibility of the gene augmentation strategy, a *Prom1^-/-^* mouse model was generated via CRISPR/Cas9 mediated deletion of exons 2–20 in the *Prom1* gene (Fig. 5A). H&E staining showed no significant difference in retinal thickness between *Prom1^-/-^* and *Prom1^+/+^* mice at postnatal week 2 (PW2). However, progressive outer retinal degeneration was observed in *Prom1^-/-^* mice from PW3 onward, consistent with the *in vivo* SD-OCT findings (Fig. 5B–E; Fig. S14BA-B). By 1 year of age, the outer retina was nearly absent in *Prom1^-/-^* mice (Fig. S14C). In contrast, *Prom^+/-^* mice show no significant changes in retinal thickness compared with *Prom1^+/+^* mice (Fig. S14D-E). Additionally, the RPE morphology of *Prom1^-/-^* mice appeared comparable to that in *Prom1^+/+^* mice (Fig. 5C). In line with the structural degeneration, functional impairment was also detected. Scotopic 3.0 fERG recordings showed marked reductions in both a- and b-wave amplitudes in *Prom1^-/-^* mice compared with *Prom^+/+^* and *Prom^+/-^* mice at postnatal month 1 (PM1) (Fig. S14G-I). It was completely abolished by 1 year of age in *Prom1^-/-^* mice (Fig. S14J). These structural and functional abnormalities demonstrate that the *Prom1^-/-^* mice faithfully recapitulate key pathological features of PROM1-IRD.

**Figure 5.**
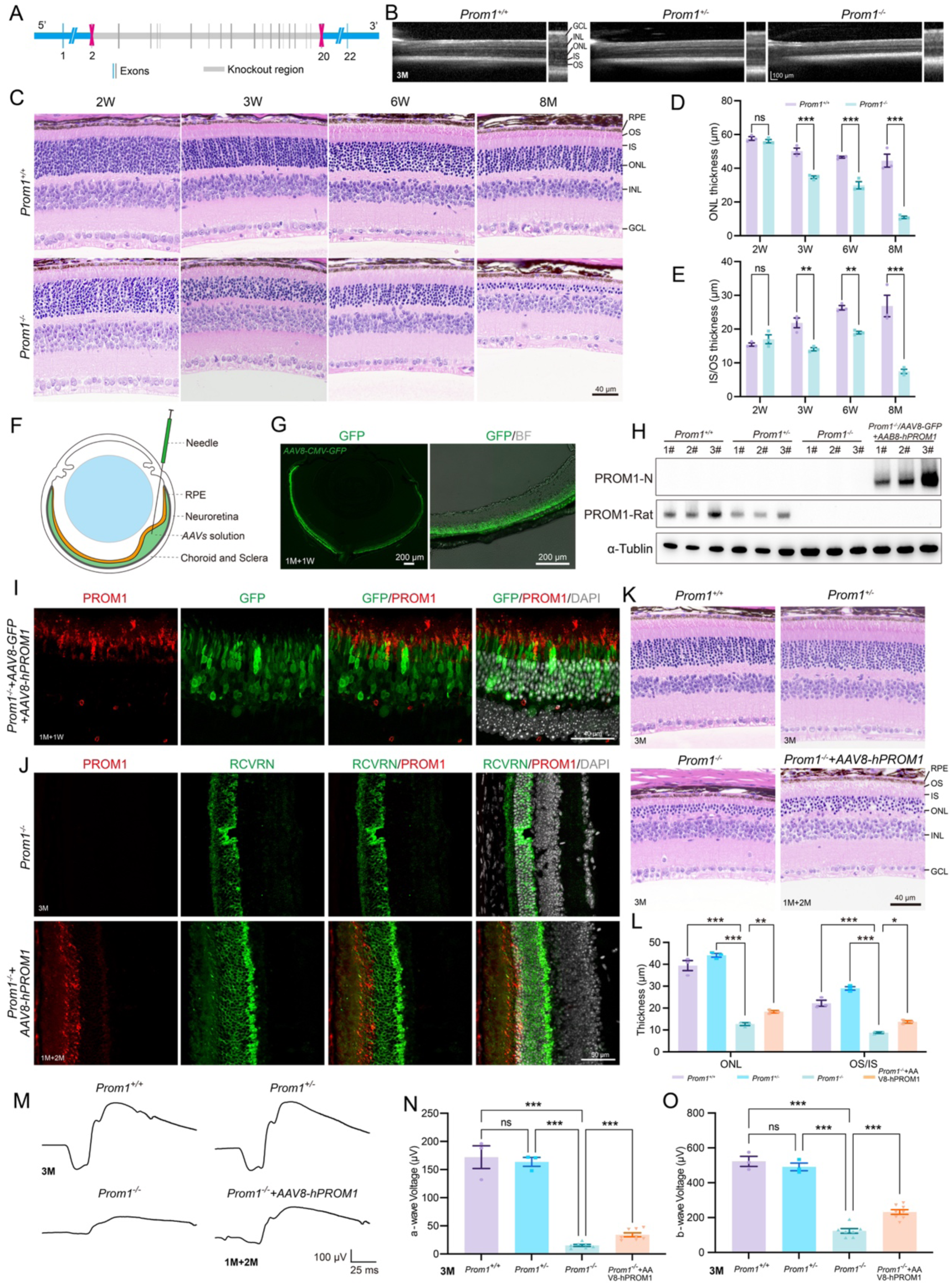
Photoreceptor-specific PROM1 expression rescues retinal degeneration in *Prom1^-/-^* mice. **(A)** Schematic of *Prom1* gene knockout strategy in mice. **(B)** SD-OCT images of 3-month-old mice retinas. Layers indicated: GCL, INL, ONL, IS and OS. Scale bar, 100 µm. **(C–E)** H&E staining of *Prom1^+/+^*, *Prom1^+/-^*, and *Prom1^-/-^* retinas. Quantification of ONL and IS/OS thickness in (D) and (E), respectively. n=3 per group. (F) Schematic of subretinal AAV injection. **(G)** Bright-field and GFP images of retina one-week post-injection of *AAV8-CMV-GFP* in *Prom1^-/-^* mice. Scale bar, 200 µm. **(H)** Western blot analysis of exogenous human PROM1 and endogenous mouse PROM1 expression in retinas. Three biological replicates labeled 1#, 2#, and 3#. **(I)** Immunostaining showing distribution of hPROM1 and GFP-labeled photoreceptors in *Prom1^-/-^* retina post co-injection of *AAV8-CRXp-hPROM1* and *AAV8-CMV-GFP* (1:10). Scale bar, 40 µm. **(J)** Immunostaining of PROM1 in the *prom1^-/-^* mice with or without *AAV8-CRXp-hPROM1* infection. Scale bar, 50 µm. **(K and L)** H&E staining and quantification of ONL and IS/OS thickness 2 months after AAV injection (n=3). Scale bar, 40 µm. **(M–O)** fERG recordings showing representative Scoptoic 3.0 waveforms (M) and quantification of a- and b-wave amplitudes (N and O). A stratified statistical analysis was performed to specifically assess gene therapy efficacy. Scale bar, 100 µV, 25 ms. Groups: *Prom1^+/+^*(n=3), *Prom1^+/-^* (n=3), *Prom1^-/-^* (n=7), and *Prom1^-/-^+AAV8-hPROM1* (n=8). *Error bars represent mean ± S.E.M.; ns, not significant (P ≥ 0.05); *P < 0.05; **P < 0.01; ***P < 0.001*.

To assess the feasibility of AAV-mediated gene therapy in this model, a mixture of *AAV8-CRXp-hPROM1* (3×10¹⁰ vg/eye) and *AAV8-CMV-GFP* (3×10⁹ vg/eye) was injected into the subretinal space of *Prom1^-/-^* mice (treated group). One week after injection, significant GFP expression was observed across the outer retina, indicating effective transduction (Fig. 5F-G). Immunoblotting with the human specific PROM1-N antibody confirmed exogenous hPROM1 expression in treated *Prom1^-/-^* retinas, with no signal in *Prom1^+/+^*, *Prom^+/-^*, or untreated *Prom1^-/-^* retinas. In contrast, the PROM1-Rat antibody detected endogenous mouse PROM1 only in *Prom1^+/+^* and *Prom^+/-^* retinas (Fig. 5H; Fig. S15). Immunostaining further revealed that hPROM1 specifically localized to the outer segments of GFP^+^ photoreceptors in treated retinas (Fig. 5I), indicating that the modified human *CRX* promoter effectively drives selective expression of hPROM1 in mice retina.

Furthermore, the therapeutic impacts of hPROM1 expression on structure and function of photoreceptors were evaluated. Two months after *AAV8-CRXp-hPROM1* (3×10¹⁰ vg/eye) injection, immunostaining confirmed sustained hPROM1 expression in photoreceptors of treated *Prom1^-/-^* mice. Immunostaining for RCVRN combined with DAPI nuclear labeling revealed a thicker photoreceptor layer in treated mice compared with untreated controls (Fig. 5J, Fig. S14F). H&E staining corroborated these findings, showing increased ONL and IS/OS thickness in treated mice (Fig. 5K-L). Functional analysis showed that, at one month post-injection, b-wave amplitudes of scotopic 3.0 fERG responses were elevated in treated mice, while a-wave amplitudes remained unchanged (Fig. S14K–M). By two months post-injection, both a- and b-wave amplitudes were significantly evaluated in treated mice relative to untreated *Prom1^-/-^* mice (Fig. 5M–O). Collectively, these results demonstrate that *AAV*-mediated delivery of human *PROM1* rescues the impairment of photoreceptor outer segments and visual function in *Prom1^-/-^* mice.

## Discussion

This study systematically characterized the expression dynamics and subcellular localization of PROM1 during photoreceptor development, revealing its photoreceptor-specific RNA alternative splicing and protein N-glycosylation patterns. To elucidate the pathogenic mechanism of PROM1-IRD, hiPSCs and ROs were generated from a patient with a *PROM1* c.619G>T homozygous mutation and his unaffected heterozygous mother. Functional analyses demonstrated that the mutation induces pronounced NMD and NAS, leading to complete loss of PROM1 protein and severe disruption of OS-like structures in Hom-ROs. Targeted gene augmentation using *AAV7m8-CRXp-hPROM1* selectively restored PROM1 expression in photoreceptors, effectively rescuing the OS-like structures. Furthermore, *in vivo* validation using *Prom1^-/-^* mice showed that subretinal delivery of *AAV8-CRXp-hPROM1* significantly slowed outer retinal degeneration and preserved visual function. Together, these findings establish a clear mechanistic link between *PROM1* mutation and photoreceptors pathology, and demonstrate the feasibility of gene augmentation as a therapeutic approach for PROM1-IRD.

PROM1 is broadly expressed in various tissues and stem cells (1–3). Although its retinal expression has been mentioned, its subcellular localization during retinal development remained unclear (12, 27, 41, 42). In this study, PROM1 expression was consistently observed throughout the entire differentiation process of ROs. As early as in RPCs, PROM1 was co-localized with ARL13B at the apical ciliary, and this distribution pattern was maintained during photoreceptor development. Although apical-basal polarity is essential for the formation and function of epithelial tissues (43), no spatial disorganization of retinal cells was observed in PROM1-deficient Hom-ROs. Photoreceptors remained properly aligned in the outer layer, indicating that loss of PROM1 does not overtly impair cell polarity during retinal differentiation. In mature photoreceptors, optimized whole-mount immunostaining revealed that PROM1 co-localized with OS markers PRPH2 and various opsins within OS-like structures. The persistent expression and precise subcellular localization of PROM1 throughout retinal development suggest its involvement in photoreceptor development, particularly in OS morphogenesis.

PROM1 exhibits a distinctive membrane topology with five transmembrane domains and two large extracellular loops that harbor nine N-glycosylation sites(7, 16). In this study, PROM1 from photoreceptors showed a substantially lower molecular weight than that in UCs, hiPSCs and hiPSC-RPE cells. TA cloning followed by Sanger sequencing revealed that *PROM1* transcripts in C1-ROs photoreceptors predominantly retained exon 4 while skipping exon 26 and/or exon 25. These isoforms were undetectable in C1-hiPSCs and align with previously reported photoreceptor-specific alternative splicing patterns (9, 32). Notably, the peptide encoded by exon 4, PETVILGLK, is proposed to function as a spacer that facilitates PROM1-ganglioside interaction, thereby contributing to OS disc morphogenesis (9, 44).

Despite the alternative splicing of exon 4 (27 bp), exon 25 (24 bp), and exon 26 (60 bp), it appeared insufficient to explain the substantial molecular weight reduction observed in photoreceptor-derived PROM1. PNGase F treatment eliminated this difference, indicating that the observed size reduction primarily resulted from photoreceptor-specific N-glycosylation. The N-Glycosylation status is closely associated with PROM1 function. For example, the high-mannose form of PROM1 facilitates its interaction with DNMT1, thereby maintaining the slow-cycling phenotype and tumorigenic capacity of glioma stem cells (17). Additionally, mutation at the N-glycosylation site Asn548 in PROM1 reduced its interaction with β-catenin and inhibited downstream β-catenin signaling (16). Our data suggest that PROM1 undergoes photoreceptor-specific post-transcriptional and post-translational modifications essential for its function in OS morphogenesis. These modifications provide a mechanistic basis for the selective vulnerability of photoreceptors to PROM1 mutations despite its broad tissue expression.

PROM1-IRDs exhibit considerable clinical heterogeneity. The age at onset, presenting symptoms, and disease severity vary depending on the specific pathogenic variants (10, 35, 45). In this study, the patient with a homozygous *PROM1* c.619C>T mutation experienced severe vision loss by age nine, indicating early and profound photoreceptor degeneration. This phenotype was faithfully recapitulated in Hom-ROs, which displayed a reduced number and shortened length of OS-like structures by day 120. Similarly, previous studies have also reported photoreceptors dysfunction as early as P12 in *Prom1^-/-^* mice (25). Given the sustained expression of PROM1 throughout photoreceptor development, we speculate that retinal damage in PROM1-IRD patients may begin during embryonic stages. Therefore, earlier therapeutic intervention may yield better outcomes in PROM1-IRD.

*PROM1* mutations can be inherited in either autosomal recessive or dominant patterns (10, 35). In our study family, four individuals carrying the *PROM1* heterozygous mutation exhibited normal vision and fundus appearance. Consistently, Het-ROs generated from the patient’s mother formed OS-like structures similar to controls, despite reduced PROM1 expression. This indicates a recessive mode of inheritance, where partial expression of wild-type PROM1 is sufficient to maintain photoreceptor function. Therefore, translational strategies such as readthrough therapy (such as NB84 or PTC124), which promote ribosomal readthrough of premature stop codons and partially restore full-length protein production, might be considered (46, 47). However, the *PROM1* c.619C>T mutation activated NMD and NAS, leading to a marked reduction in mRNA levels. Thus, readthrough therapies are unlikely to be effective for this PROM1-IRD. Notably, nonsense mutations account for approximately 18.5% of IRD-related variants (48). These mutations often elicit RNA surveillance mechanisms beyond truncated protein production (36, 37). Therefore, assessing the impact of nonsense mutations at both transcript and protein levels is essential before considering readthrough therapy for IRDs.

*AAV*-medicated gene augmentation therapy has emerged as a promising treatment for IRDs (19). Despite *PROM1* being a relatively frequent causative gene for IRDs, no such efforts have yet been reported (24). Our study fills this gap by systematically evaluating the feasibility of PROM1 gene augmentation. Patient-derived ROs offer an ideal platform for preclinical testing, as they accurately mimic native retinal development and disease (30, 31). However, variability in retinal differentiation protocols has resulted in inconsistent *AAVs* transduction time points, ranging from D40 to beyond D200 (49–51). Previous studies, including those based on our earlier protocols (30), transduced *AAV*s at late stages (≥D120) (50, 52). Given the early and continuous expression of PROM1, we aimed to identify earlier transduction windows. At D60, a timepoint when RCVRN⁺ photoreceptors are just emerging (31), *AAVs* transduction caused severe toxicity and cell death. In contrast, D90 infection, prior to OS-like structure formation (PRPH2⁻), caused no detectable damage. These findings identify D90 or later as an optimal window for *AAVs* transduction in ROs.

We further optimized gene delivery in ROs by selecting an efficient promoter and *AAV* serotype. A modified *CRX* promoter was chosen to drive photoreceptor-selective expression of hPROM1 (53). *CRX* is a key transcription factor that governs the expression of the majority of rods- and cones-specific genes and is indispensable for photoreceptor development (54). Our previous work confirmed that CRX expression persists during photoreceptor development and maturation in ROs (31), providing strong rationale for the use of its promoter. Among the *AAV* serotypes screened, *AAV7m8-CMV-GFP* exhibited the highest transduction efficiency in D90 ROs, outperforming *AAV2* and *AAV8*. Additionally, *AAV7m8* has also been widely used in prior IRD gene therapy studies involving ROs (55–57). Based on these, we selected *AAV7m8-CRXp-hPROM1* to transduce Hom-ROs. One month post-transduction, PROM1 expression was restored in over half of photoreceptors, significantly preserved OS-like structures, underscoring the therapeutic potential of this approach in PROM1-IRD.

To further evaluate the therapeutic efficacy of this strategy, we generated a *Prom1^-/-^* mouse model. By one year of age, this model exhibited severe outer retinal thinning and extinguished fERG response, in line with prior studies (25–28). However, ONL and IS/OS thicknesses remained comparable to wild-type at PW2, and degeneration progressed gradually until PM3, with partial preservation of visual function. Except for LCA, many IRDs are characterized by a slow progression of visual decline and photoreceptor degeneration, with a substantial number not experiencing severe functional impairment until adulthood (48). Therefore, compared with previously reported *Prom1^-/-^* mice exhibiting rapid photoreceptor loss (26), the model established in this study more closely recapitulates the slow-progressive degeneration phenotype of PROM1-IRD. For IRD therapeutic research, a slower retinal degenerating model permits broader intervention windows and simplifies surgical procedures post-eyelid opening. Thus, we believe the *Prom1^-/-^* mice developed here is well suited for studying both IRDs pathogenesis and treatment.

Using this model, we validated that *AAV8-CRXp-hPROM1* efficiently restored hPROM1 expression in photoreceptors, with selective localization to the OS. Notably, the PROM1-N antibody does not recognize endogenous mouse PROM1, whereas the PROM1-Rat antibody does. It appears due to the limited (∼60%) homology between primate and rodent PROM1 proteins (8, 58). Despite these species difference, human PROM1 expression significantly ameliorated photoreceptor disruption and preserved visual function in *Prom1^-/-^* mice, confirming the therapeutic potential of this gene augmentation strategy.

In summary, this study elucidates the role of PROM1 in photoreceptor development and reveals the pathogenic mechanisms underlying PROM1-IRD. More importantly, it provides the first preclinical proof-of-concept for AAV-mediated gene augmentation therapy in both patient-derived retinal organoids and *Prom1⁻/⁻* mice. These findings not only support the clinical translation of gene therapy for PROM1-IRD, but also offer a conceptual framework for investigating and treating related forms of inherited retinal degeneration.

## Methods

### Study Design

This study aimed to elucidate the molecular basis of PROM1-IRD and develop a feasible gene augmentation strategy. PROM1 expression patterns in photoreceptors were examined using wild-type UCs, hiPSCs, hiPSC-RPE cells, and ROs. To investigate disease mechanisms, patient-specific hiPSCs carrying the homozygous c.619G>T (p.E207X) mutation and a heterozygous parental control were differentiated into ROs, and the effects of the mutation on mRNA splicing, protein expression, and photoreceptor morphology were systematically analyzed. To evaluate therapeutic potential, photoreceptor-specific *AAV7m8-CRXp-hPROM1* was used to infect patient-derived ROs at optimized developmental stages, and the restoration of PROM1 expression and OS-like structures was assessed by whole-mount immunofluorescence and morphometric analysis. For *in vivo* validation, a *Prom1⁻/⁻* mouse model was generated via CRISPR/Cas9-mediated deletion of exons 2–20, followed by subretinal injection of *AAV8-CRXp-hPROM1*. Retinal structure and visual function were evaluated by histology, immunostaining, and fERG. Approximately equal numbers of male and female mice were used in each experiment to assess potential sex-specific effects. Investigators were not blinded to sample allocation, data acquisition, or data analysis (except for image quantification). Mice with severe endophthalmitis or corneal neovascularization were excluded. All experiments were conducted with appropriate controls and replicates, and statistical analyses were performed as detailed in the Methods.

### Ethics approval

All human studies were conducted in accordance with the Declaration of Helsinki and approved by the Ethics Committee of the Zhongshan Ophthalmic Center, Sun Yat-sen University (Approval No. 2017KYPJ061). Written informed consent was obtained from all participants. All animal procedures were approved by the Institutional Animal Care and Use Committee (IACUC) of the Zhongshan Ophthalmic Center, Sun Yat-sen University (Approval No. OW2022003), and were conducted in accordance with institutional and national guidelines for animal care and use.

### Urine cells and hiPSC lines

Four hiPSC lines were used in this study. UCs were collected from midstream urine samples of a patient harboring a homozygous nonsense mutation in *PROM1* [c.619G>T (p.E207X)] and his mother. These UCs were reprogrammed into hiPSCs using previously reported protocols (38). The patient-specific hiPSC line was designated SKLOi002-A (Hom-hiPSCs), and the maternal line was designated SKLOi003-A (Het-hiPSCs). Two additional control hiPSC lines (C1 and C2) were generated from the urine cells of a healthy volunteer. All hiPSC lines were cultured on Matrigel-coated plates (Corning, 354277) in mTeSR1 medium (STEMCELL Technologies, 05851) and passaged as clumps at a 1:8 ratio every 4–6 days.

### Differentiation of hiPSCs into ROs

HiPSCs were differentiated into ROs using a modified version of our previously established protocol (30, 59). Briefly, hiPSCs at >80% confluence were collected for suspension culture to allow embryo body (EB) formation. Over the next 3 days, the medium was gradually switched to NIM. From D4 to D6, EBs exhibiting translucency and appropriate morphology were plated onto Matrigel-coated dishes. On D16, the medium was replaced with RDM. During weeks 4– 5, optic vesicle-like structures were manually isolated transferred to suspension culture in RDM. From week 6 onward, cultures were maintained in RC1 medium, which was subsequently replaced with RC2 medium after week 12 to support long-term maturation.

### *AAV* vectors generation and virus production

Recombinant plasmids containing a modified human *CRX* promoter were utilized to drive the exogenous expression of human PROM1 protein in photoreceptors. The modified *CRX* promoter spans 631 bp and is composed of three segments derived from the 5′ untranslated region of the human *CRX* gene (NG_008605.1) (53). Plasmid construction and AAV packaging were performed by VectorBuilder (Guangdong, China) using a standard triple transfection protocol in HEK293 cells. Three AAV serotypes were generated, including *AAV2-CRXp-hPROM1*, *AAV7m8-CRXp-hPROM1*, and *AAV8-CRXp-hPROM1*. Corresponding CMV-GFP control vectors were also produced. All preparations had viral titers exceeding 1.0 × 10¹³ vg/mL.

### *AAVs* infection of ROs

ROs were transferred to a 48-well culture plate, with one RO per well. *AAVs* were added at the appropriate concentration based on the different experimental requirements in this study. After 48 hours of infection, the culture medium was aspirated, and the ROs were washed three times with DMEM basic. Subsequently, the ROs were transferred to a 24-well plate, and stage-specific ROs culture medium was added for continued culture. The status of the ROs was monitored and recorded using a phase-contrast microscope (Nikon) or an Axio inverted fluorescence microscope (Zeiss).

### Whole-mount immunostaining and imaging of ROs

ROs were fixed in 4% PFA (DF0135, Leagene Biotechnology) at room temperature for 30 minutes, washed in PBS, and permeabilized with 0.225% Triton X-100 in 10% donkey serum (Ruite, W9030-01) for 30 minutes. Then, ROs were incubated overnight at 4°C with primary antibodies, followed by PBS washes and secondary antibody incubation at room temperature for 1 hour (Table S1). Nuclei were counterstained with DAPI (Dojindo). ROs were mounted in antifade medium (Beyotime, P0126) and imaged using an inverted fluorescence microscopy or a Zeiss LSM 880 confocal microscope (Zeiss). The lengths of OS-like structure length were quantified using ImageJ (NIH).

### Immunofluorescent staining

Cells on coverslips or Transwell membranes were fixed with 4% PFA for 10 minutes at room temperature. ROs were fixed for 30 minutes, while mouse eyes were fixed overnight at 4°C. Fixed tissues were dehydrated in graded sucrose (6.25%, 12.5%, and 25%), embedded in OCT (Sakura, OCT4583), and sectioned at 12 μm. Sections or cell samples were stained with primary antibodies overnight at 4°C, followed by secondary antibody incubation. All antibodies used are listed in Table S1. Imaging was performed using a Zeiss LSM 880, Axio Scan Z1, or inverted microscope, and processed with Zen software.

### Western blot and PROM1 glycosylation analysis

Cell samples were lysed with ice-cold RIPA buffer containing 1 mM PMSF (ST506, Beyotime) for 15 minutes. ROs and mouse retinas were subjected to ultrasonic treatment. All samples were then centrifuged to collect supernatants. For PROM1 N-glycosylation analysis, supernatants were treated with PNGase F (4450, Takara) according to the manufacturer’s protocol. Proteins were resolved via SDS-PAGE, transferred to PVDF membranes, and probed with primary and secondary antibodies (Table S1). Detection was performed using chemiluminescence (P0018FS, Beyotime), and band intensity quantified with ImageJ.

### TA Cloning and sequencing

*PROM1* cDNA fragments were PCR-amplified from hiPSCs or ROs and gel-purified (QIAGEN, 281006). Products were ligated into the *pMD19-T* vector (Takara, 3271) and transformed into DH5α E. coli (Solarbio, C1100). Positive colonies were selected for plasmid extraction (Magen, P1002) and subjected to Sanger sequencing. Primer sequences are listed in Table S2.

### Murine subretinal injection

A 2057 bp genomic deletion encompassing exons 2–20 of the *Prom1* gene (NM_001163582) was introduced into C57BL/6JCya mouse via CRISPR/Cas9 (Cyagen, Guangzhou). Littermate Prom1^+/+^ and Prom1^+/-^ mice were used as controls. For subretinal injection, mice were anesthetized with 1% sodium pentobarbital. A scleral incision was created approximately 1 mm posterior to the limbus using a 10-0 suture needle under a surgical microscope. A 2.5 µL Hamilton syringe fitted with a 33G needle was inserted through the incision, taking care to avoid the lens. A total of 1 µL of *AAVs* solution was injected into the subretinal space. Successful injection was indicated by localized retinal detachment and vascular bending. Tobramycin ophthalmic ointment was applied to the operated eye.

### Retinal Electrophysiology and Imaging in Mice

fERG was recorded using the RETI-port/Scan 21 system (Roland, Germany) after ≥8 hours of dark adaptation. Mice were anesthetized and pupils dilated (tropicamide), with corneal anesthesia (proparacaine) and lubrication (2% HPMC). A reference electrode was placed subcutaneously near the orbit, with the active electrode on the cornea and the ground electrode on the tail. fERG responses included scotopic 0.01, 3.0, and 10.0 ERGs. SD-OCT imaging was performed using the Spectralis HRA+OCT system (Heidelberg Engineering, Germany) in high-speed mode, with each scan averaged over 12 frames. Images were analyzed using Heidelberg Eye Explorer software.

### Statistical analysis

Statistical analyses for RNA-seq data are detailed above. Other statistical tests were performed using GraphPad Prism version 9.0 (GraphPad Software). Comparisons between two groups were conducted using two-tailed Student’s t-tests; comparisons among three or more groups were analyzed using one-way or two-way ANOVA, as appropriate. Specially the *Prom1^−/−^* and *Prom1^−/−^+AAV8-hPROM1* groups exhibited markedly lower values than the *Prom1^+/+^* and *Prom1^+/−^* groups (Figure 5N), potentially obscuring treatment effects. To resolve this, a stratified analysis was performed. Firstly, a one-way ANOVA was used to compare the *Prom1^+/+^*, *Prom1^+/−^*, and *Prom1^−/−^* groups to assess the effect of the *PROM1* mutation. Then, to specifically evaluate the therapeutic effect of gene augmentation, a two-tailed Student’s t-test was applied to compare the *Prom1^−/−^* and *Prom1^−/−^+AAV8-hPROM1 groups*. All experiments were performed in triplicate unless otherwise stated. Data are presented as mean ± standard error of the mean (SEM). Statistical significance was defined as ns (not significant, *P ≥ 0.05*), \**P < 0.05*, *\*\*P < 0.01*, and *\*\*\*P < 0.001*.

## Data and materials availability

All data supporting the findings of this study are available in the main text or the Supplementary Materials. Raw RNA-seq data have been deposited in the NCBI GEO under accession number GSE299857. Plasmids, AAVs, and mouse models used in this study can be obtained from commercial sources approved in the manuscript. HiPSC lines are available through an MTA.

## Acknowledgments

We thank the patient and his family for their participation in this study. We are also grateful to Dr. Jeremy Nathans (Johns Hopkins University) for generously providing S-opsin and L/M-opsin antibodies. This work was supported partly by the National Key R&D Program Project of the Ministry of Science and Technology (2023YFC2506100, 2017YFA0104101), the National Natural Science Foundation of China (81970842, 82172957), Science & Technology Project of Guangdong Province (2017B020230003), a joint grant from the Science and Technology Project of Guangzhou and Zhongshan Ophthalmic Center, Sun Yat-sen University (202201020312), the China Postdoctoral Science Foundation (2023M744064), the Guangdong Basic and Applied Basic Research Foundation (2022A1515110631), and the GBRCE for Major Blinding Eye Diseases Prevention and Treatment.

## Author contributions

Conceptualization: XZ, PX, YW. Methodology: PX, FG, YW. Investigation: PX, FG, YW, XS, BYL, DZ, GG, WY, SZ. Visualization: PX, YW, FG. Funding acquisition: XZ, PX, GG. Project administration: XZ, PX, BTL. Supervision: XZ. Writing – original draft: XZ, PX, FG, YW. Writing – review & editing: XZ, PX.

## Supplement methods

### Ophthalmic examination and diagnosis

A 13-year-old male patient initially presented with blurred vision and was diagnosed with IRD at Tongji Hospital (Wuhan, China). The ophthalmic assessments were performed at ages 13 and 35, including best-corrected visual acuity (BCVA), intraocular pressure, color fundus photography, fluorescein angiography (FFA) and genetic diagnosis. At age 38, he was referred to the Zhongshan Ophthalmic Center for comprehensive ophthalmic evaluation, including spectral-domain optical coherence tomography (SD-OCT), indocyanine green angiography (ICGA), flash electroretinography (fERG), and flash visual evoked potentials (fVEP). Additionally, the patient’s mother also underwent ophthalmic assessment. Written informed consent for participation and publication was obtained from both individuals.

### Expansion and culture of hiPSC-RPE cells

Procedures were performed as previously described(*1, 2*). Briefly, after ROs isolation, the residual tissue was cultured in RC1 to generate hiPSC-RPE cells. At 7–8 weeks, pigmented hiPSC-RPE cells with cobblestone morphology were manually isolated and maintained in suspension in RPE expansion medium (50% DMEM/F12, 35% basic DMEM, 10% FBS, 100 µM taurine, 2% supplement, 1% GlutaMAX, 1% MEM NEAA, 1% AA). After 2 weeks, cells were dissociated with TrypLE Express and seeded onto Matrigel-coated plates. Upon confluence, cells were cryopreserved. For subsequent experiments, P3 cells were plated at 5×10⁴ cells/cm² on Matrigel or Transwells in expansion medium. After 5–7 days of culture, the medium was switched to RPE maturation medium (59% DMEM/F12, 37% DMEM, 2% B27, 1% FBS, 1% AA) for long term culture.

### RNA-seq and data analysis

ROs (n=6 per group) were harvested on day 150 (3 biological replicates per group). Photoreceptors layers were mechanically dissected and submitted for RNA-seq (Gene Denovo, Guangzhou). Gene expression was quantified as FPKM. Differentially expressed genes (DEGs) were identified using a false discovery rate (FDR) < 0.05 and absolute log₂(fold change) > 1. GO and KEGG enrichment analyses were performed; top 10 pathways were plotted. DEG heatmaps, histograms, and pathway visualizations were generated using Omicsmart (www.omicsmart.com).

### TUNEL assay

Frozen sections of D150 ROs were washed with PBS. Following the manufacturer’s instructions, 50 μl of TUNEL reaction mixture (Roche, 121567929101) was added to each sample and incubated at 37°C for 1 hour. ROs were counterstained with DAPI to label the nuclei. Images were acquired on a Zeiss LSM 880 confocal microscope and analyzed using Zen software.

### Electron microscopy analysis

D150 ROs were fixed in modified Karnovsky’s solution (2% glutaraldehyde and 2% paraformaldehyde) overnight at 4°C. The samples were then sent to the Electron Microscopy Core Facility of Sun Yat-sen College of Medical Science, Sun Yat-sen University (Guangzhou, China) for scanning electron microscopy (SEM) analysis. Samples were dehydrated through a graded ethanol series and subsequently dried using the Leica EM CPD 300. After drying, the samples were sputter-coated with gold using a Coater Ion sputter EIKO IB-5. SEM images were captured at an acceleration voltage of 10 kV with a lower secondary electron detector using a FEI Quanta 200 SEM (FEI, Thermo Fisher Scientific).

### RT-PCR and Sanger sequencing

Genomic DNA was extracted using a DNA Quick Extraction Kit (D0065M, Beyotime). Total RNA was extracted using Trizol reagent (T9424, Sigma-Aldrich), and reverse-transcribed into cDNA using a Reverse Transcription Kit (RR047A, Takara). Target regions were amplified by PCR using primers in Table S2. PCR products were sent to Sangon Biotech Company (China) for Sanger sequencing. Sequencing data were analyzed using SnapGene software v5.0 (SnapGene) and DNAMAN software v8.0 (Lynnon Biosoft).

### H&E Staining

Mice were euthanized by cervical dislocation; eyes were enucleated, rinsed with PBS, and fixed in 4% PFA overnight at 4 °C. Samples were processed by Servicebio (Wuhan, China) for paraffin embedding and sectioning. Sections containing the optic disc were stained with H&E, imaged on an Axio Scan Z1, and analyzed using Zen software (Zeiss).

## Supplement results

**Figure S1.**
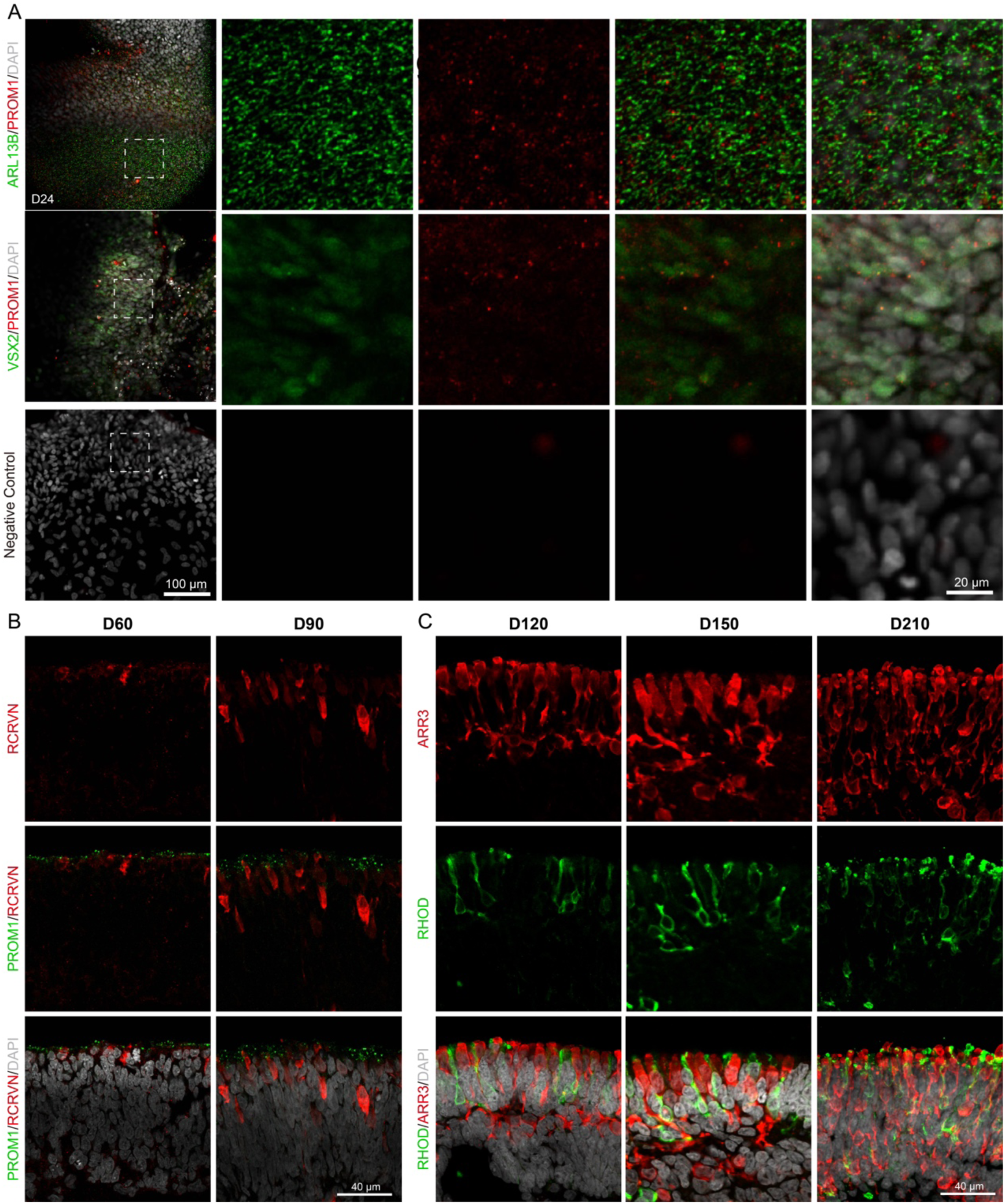
Expression and distribution of PROM1 in photoreceptors during ROs differentiation. **(A)** Immunostaining of PROM1, ARL13B, and VSX2 in optic-cup-like structures at D24 of C1-ROs differentiation. Antibody specificity was confirmed using negative controls. Magnified regions are indicated by white dashed boxes. Scale bars: 100 µm (overview) and 20 µm (magnified view). **(B)** Immunostaining of RCVRN and PROM1 in C1-ROs at D60 and D90. Scale bar, 40 μm. **(C)** Expression and distribution of rod (RHOD^+^) and cone (ARR3^+^) in C1-ROs at D120, D150, and D210. Scale bar, 40 μm.

**Figure S2.**
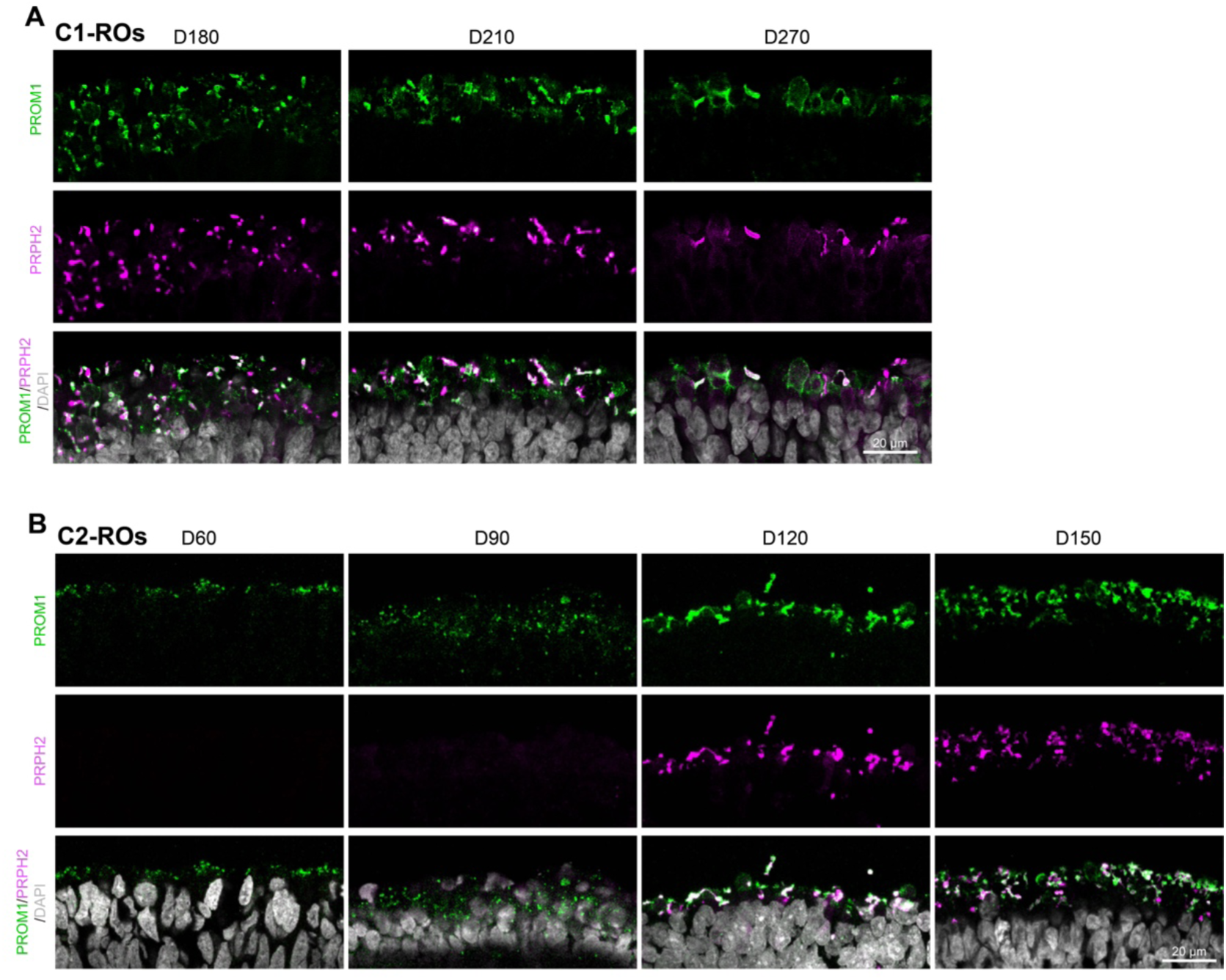
Expression of PROM1 in OS-like structures in ROs. Immunostaining shows the expression and distribution of PROM1 in both ROs at multiple time points. PRPH2 was used to label OS-like structures. C1-ROs and C2-ROs were derived from two independent wild-type hiPSC lines. Scale bar, 20 μm.

**Figure S3.**
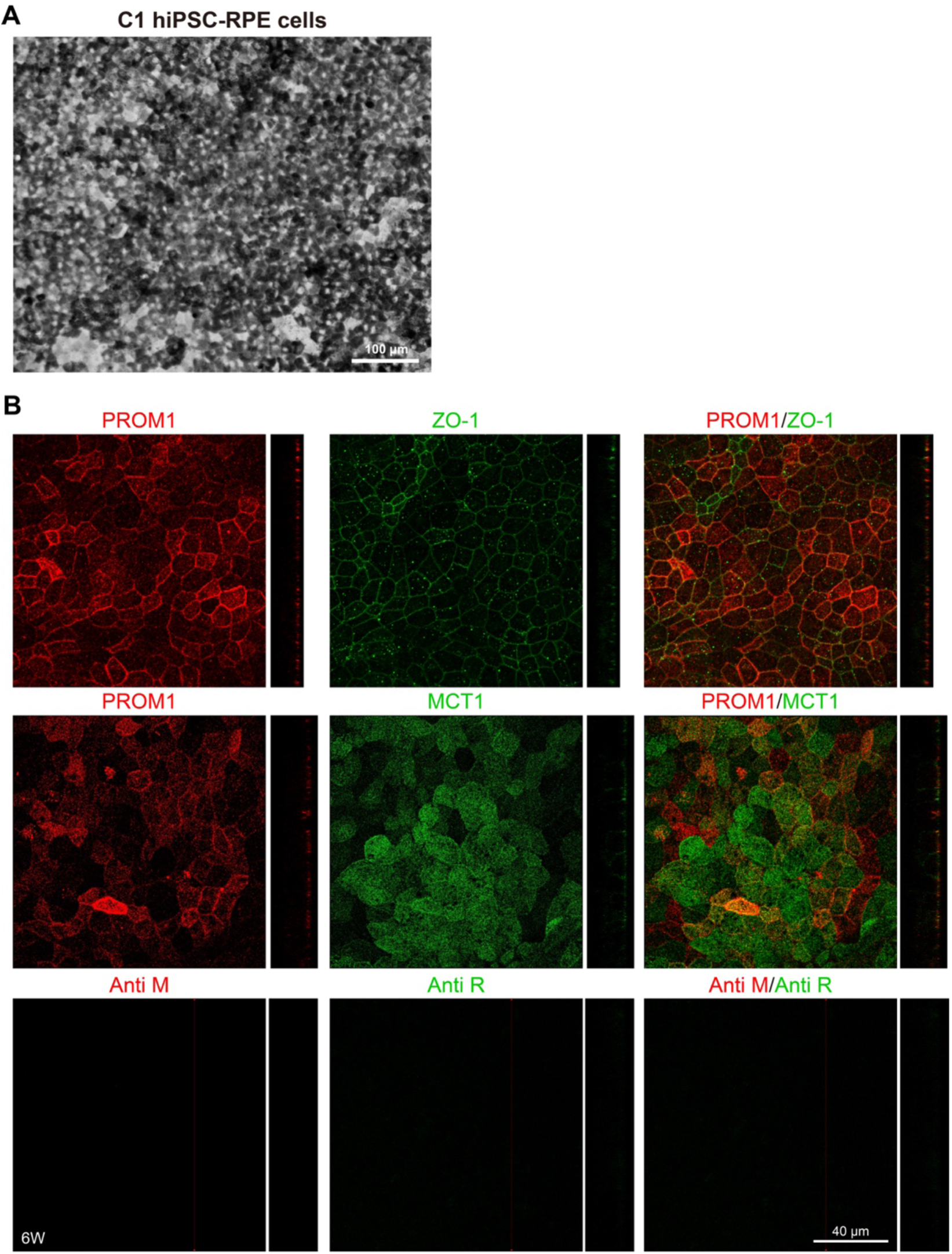
Subcellular localization of PROM1 in hiPSC-RPE Cells. **(A)** Bright-field image of C1 hiPSC-RPE cells cultured on Transwells for 6 weeks. Scale bar, 100 μm. **(B)** Immunostaining of PROM1 together with MCT1 and ZO-1 in C1 hiPSC-RPE cells, corresponding to Figure 1F and 1G. Scale bar, 40 μm. The negative control confirms specificity. Scale bar, 40 μm.

**Figure S4.**
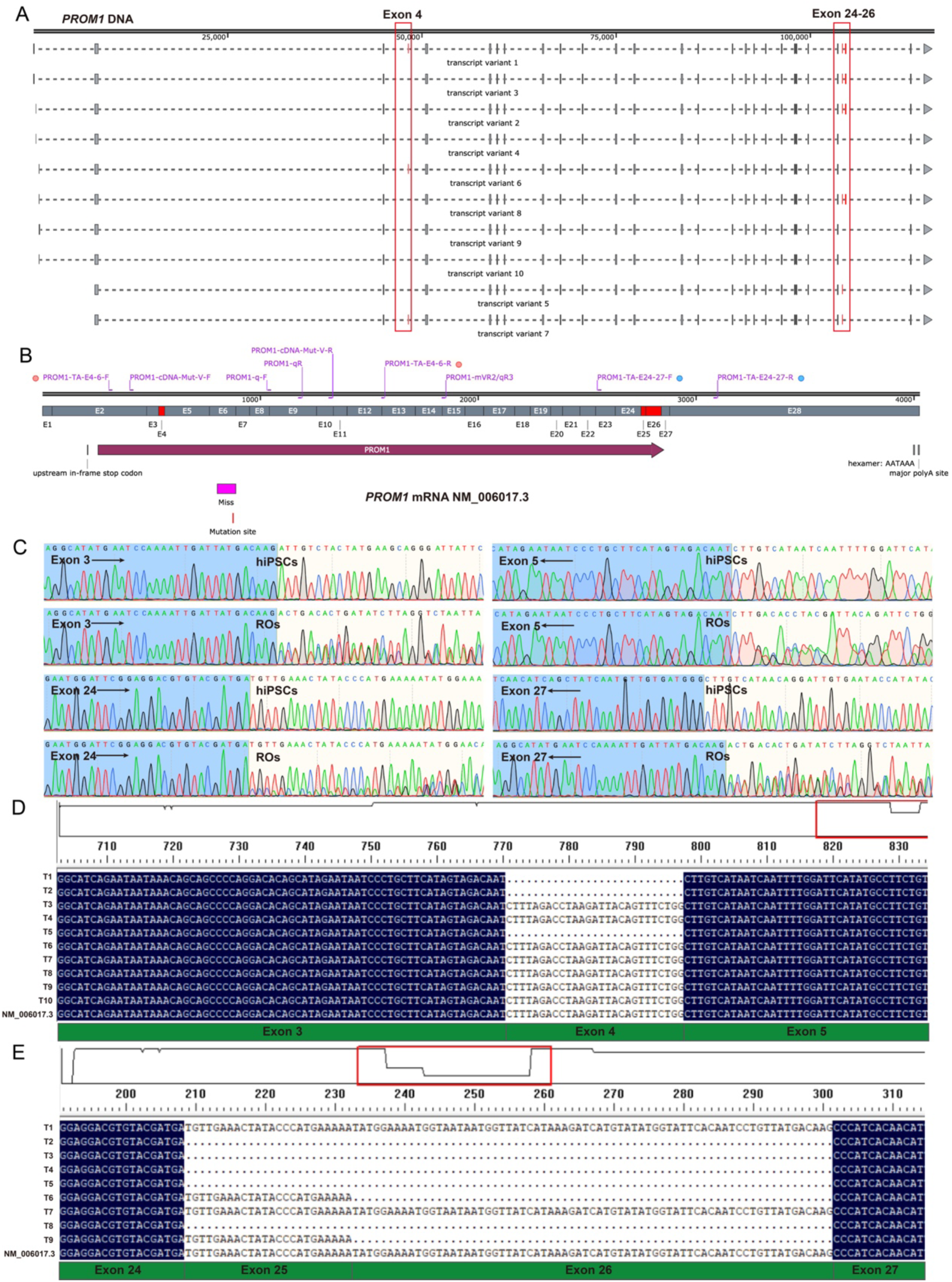
*PROM1* mRNA alternative splicing in hiPSCs and ROs. **(A)** Schematic of the *PROM1* genomic structure and ten alternatively spliced mRNA isoforms; transcript variant 1 (NM_006017.3) is the longest, containing all 28 exons. Major splicing differences at exon 4 and exons 25–26 are highlighted. **(B)** Diagram of PROM1 mRNA showing primer binding sites used for RT-PCR. **(C)** Bidirectional Sanger sequencing of PROM1 splice variants involving exon 4 and exons 25–26 in C1-hiPSCs and C1-ROs; sequencing directions are indicated by black arrows. **(D and E)** TA cloning and Sanger sequencing of PROM1 transcripts spanning exons 3–12 (D) and exons 24–27 (E) from C1-ROs.

**Figure S5.**
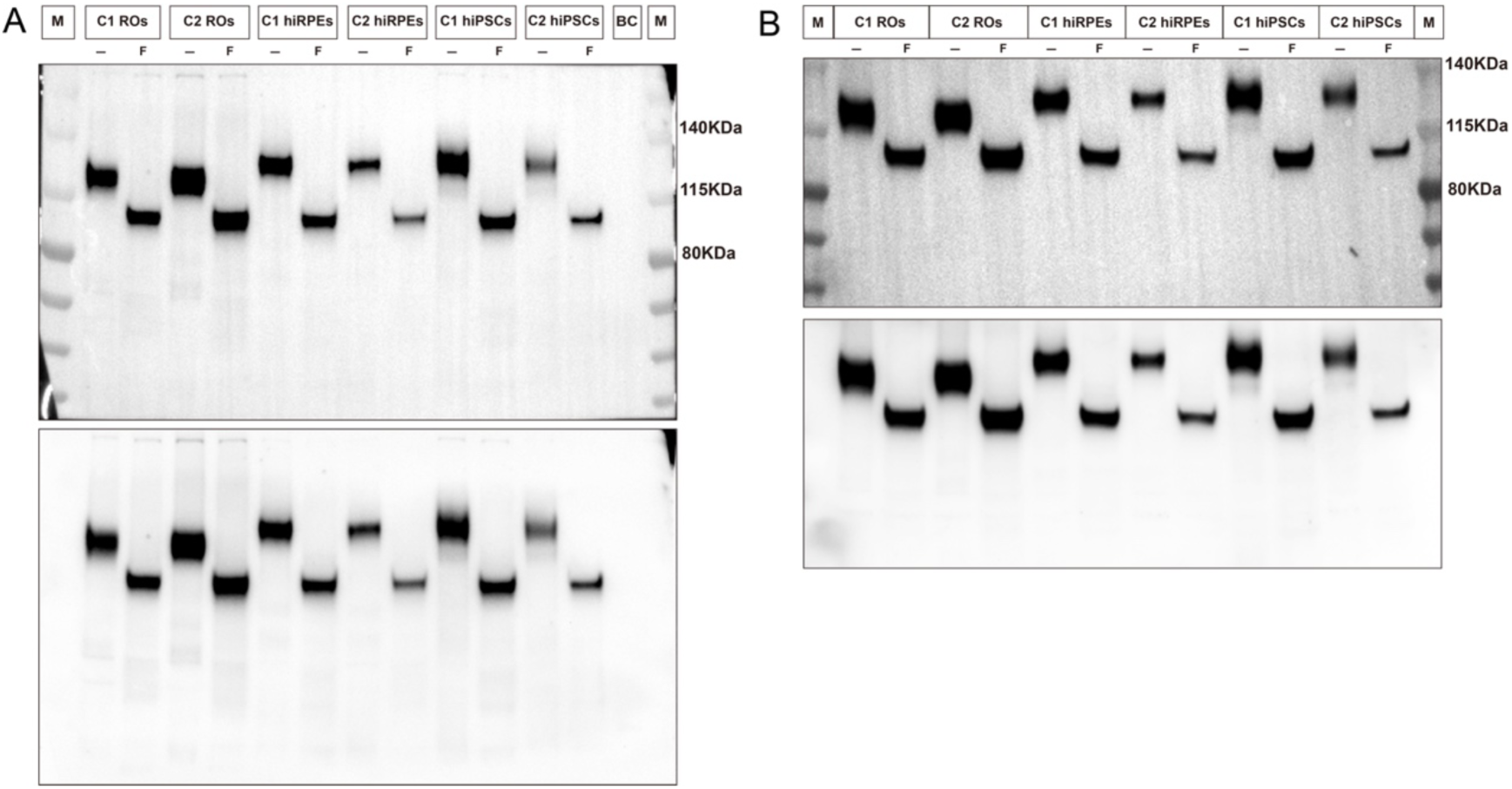
Comparative expression of PROM1 in various cell types. **(A)** Original Western blot image corresponding to Figure 1H. **(B)** Technical replicate of the same samples. “BC”, blank control; “M”, pre-stained protein marker; “F”, Glycopeptidase F-treated samples; “−”, untreated controls.

**Figure S6.**
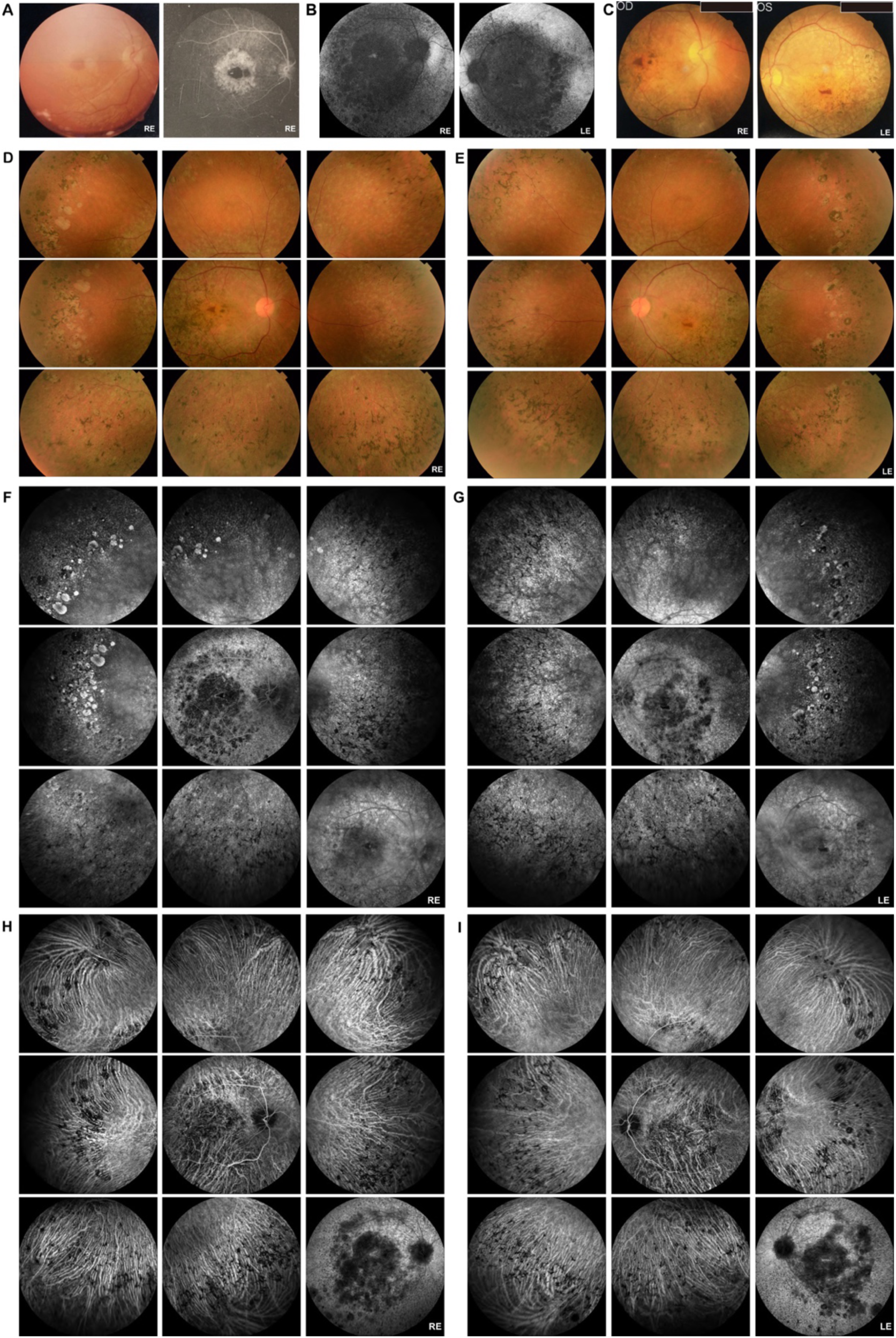
Ocular features of the PROM1-IRD patient. **(A)** Fundus examination at age 13, including right-eye color fundus photograph and FFA **(B)** BAF imaging at age 38. **(C)** Color fundus photographs at age 32; personal information is obscured by a black box. **(D-I)** Expanded multimodal retinal imaging at age 35 (as shown in Figure 2), comprising nine-field color fundus photographs (D and E), FFA (F and G), and ICGA (H and I).

**Figure S7.**
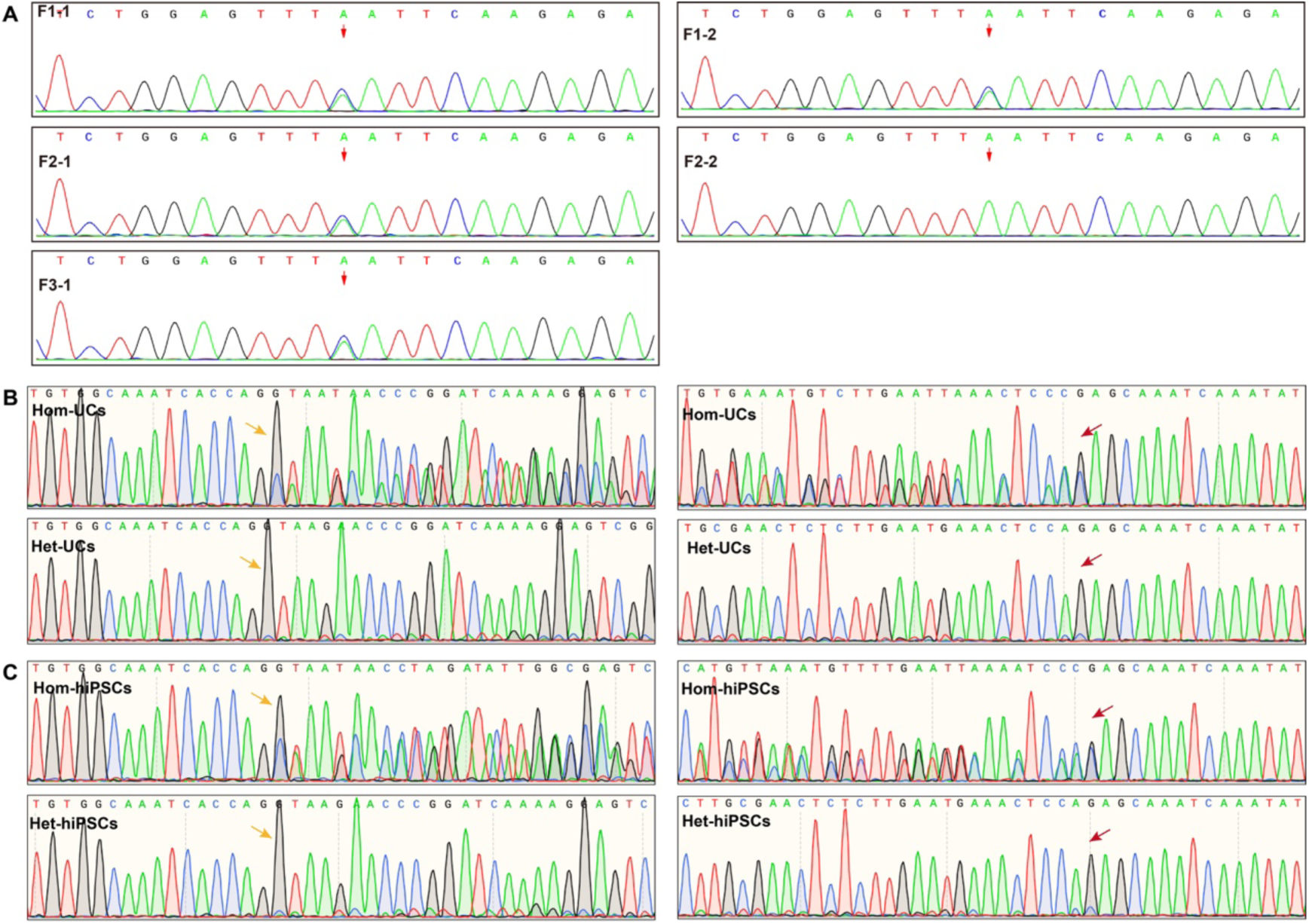
Sanger sequencing confirmed the *PROM1* c.619G>T Mutation. **(A)** Detection of the *PROM1* mutation in members of the PROM1-IRD pedigree. Red arrow indicates the mutation site. **(B and C)** Sanger sequencing analysis of the *PROM1* mutation in cDNA from UCs and hiPSCs. The left panel shows the beginning of overlapping peaks at position c.548 (yellow arrow), while the right panel indicates the disappearance of these overlaps at position c.630 (red arrow).

**Figure S8.**
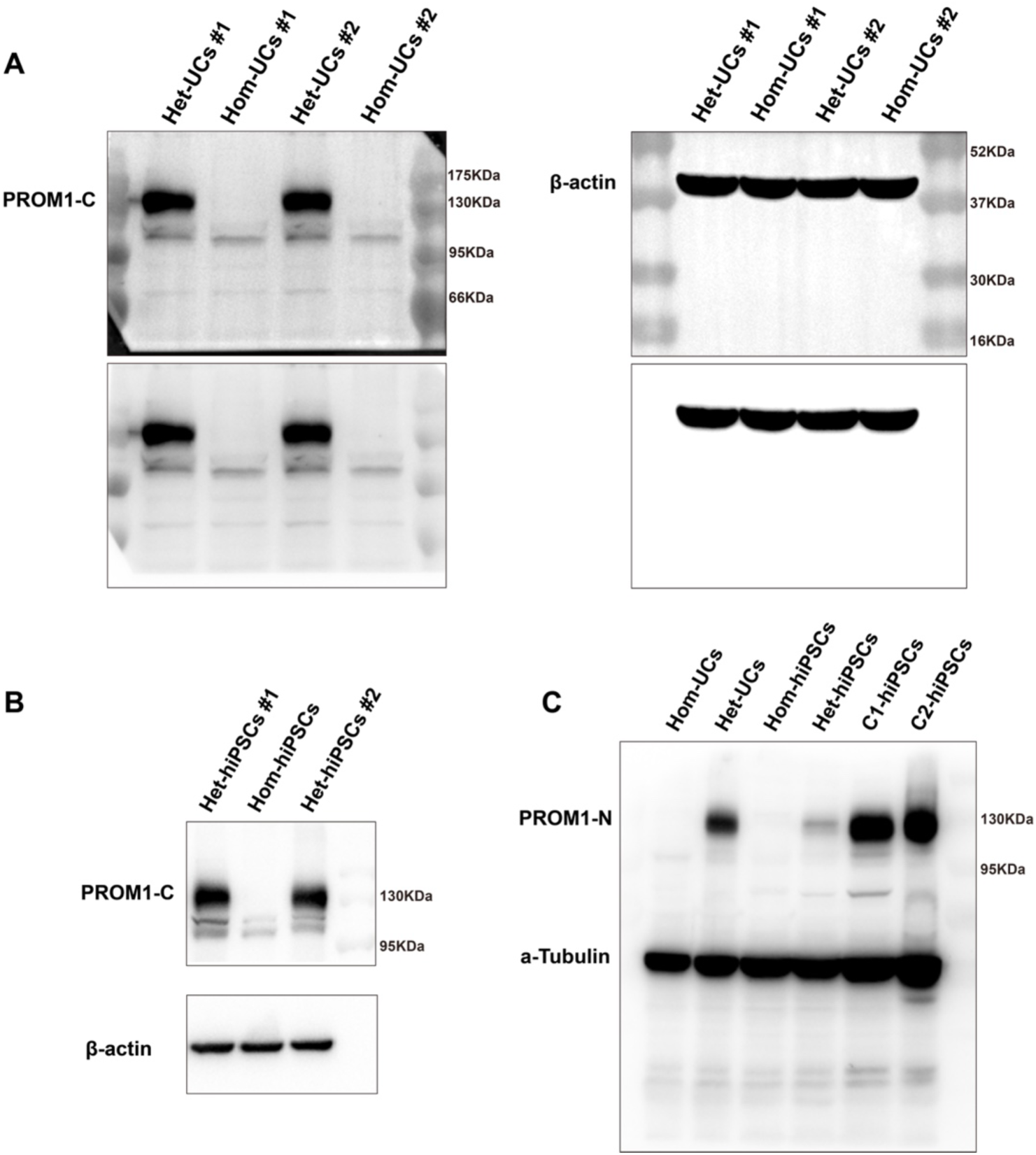
Detection of PROM1 protein using different antibodies. **(A and B)** Original Western blot images correspond to Figures 2F and 2H. **(C)** PROM1 expression in UCs and hiPSCs was assessed using the PROM1-N antibody, which specifically recognizes the N-terminal region (aa 20– 108) of PROM1. α-Tubulin serves as loading control.

**Figure S9.**
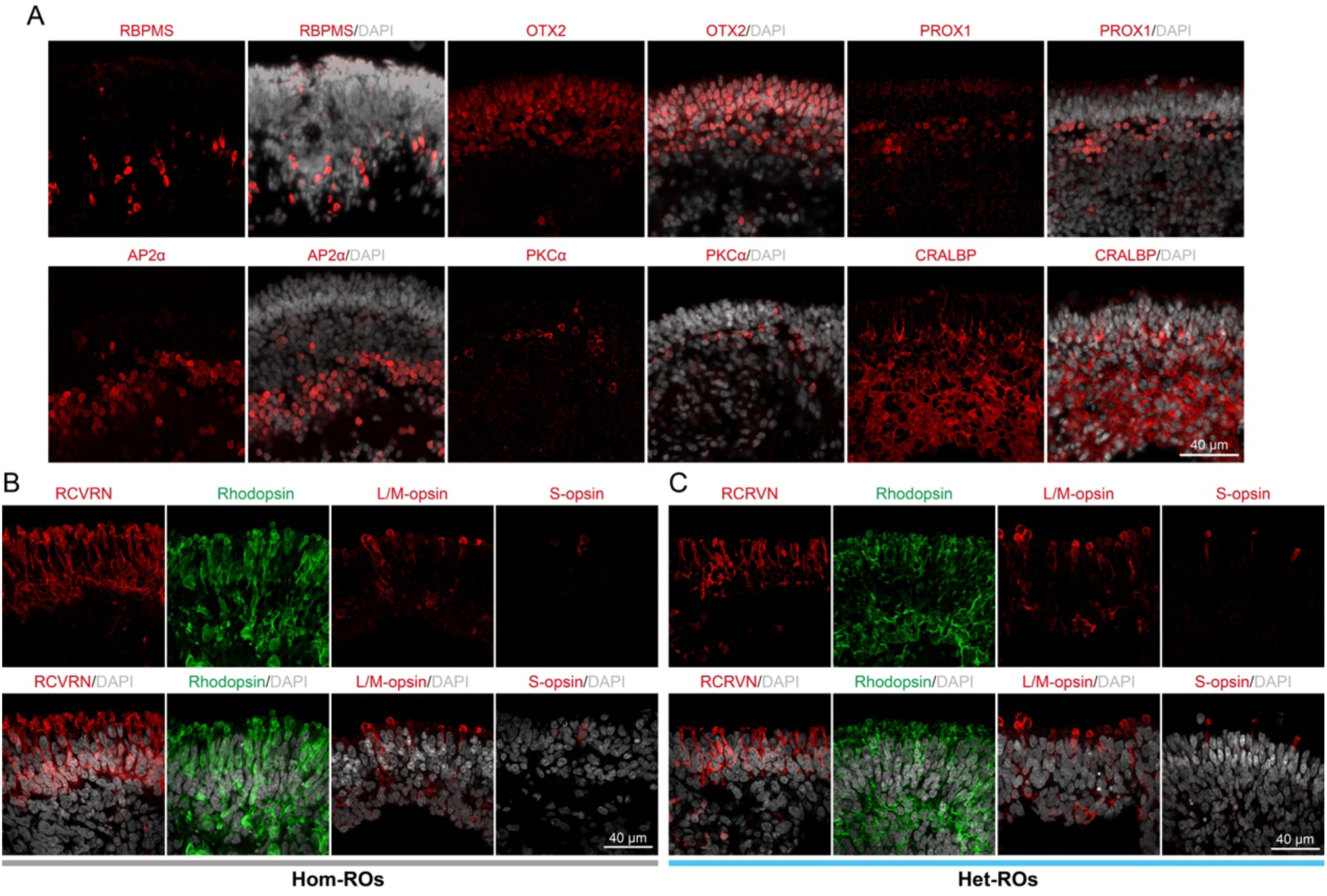
Patient-specific ROs maintain all types of neural retinal cell types. **(A)** Immunostaining of Hom-ROs demonstrates the distribution of various neural retinal cell types using markers RBPMS, OTX2, PROX1, AP2α, PKCα, and CRALBP. Scale bar, 40 μm. **(B and C)** Photoreceptors in Het-ROs and Hom-ROs were identified using markers RCVRN, Rhodopsin, L/M–opsin, and S–opsin. Scale bar, 40 µm.

**Figure S10.**
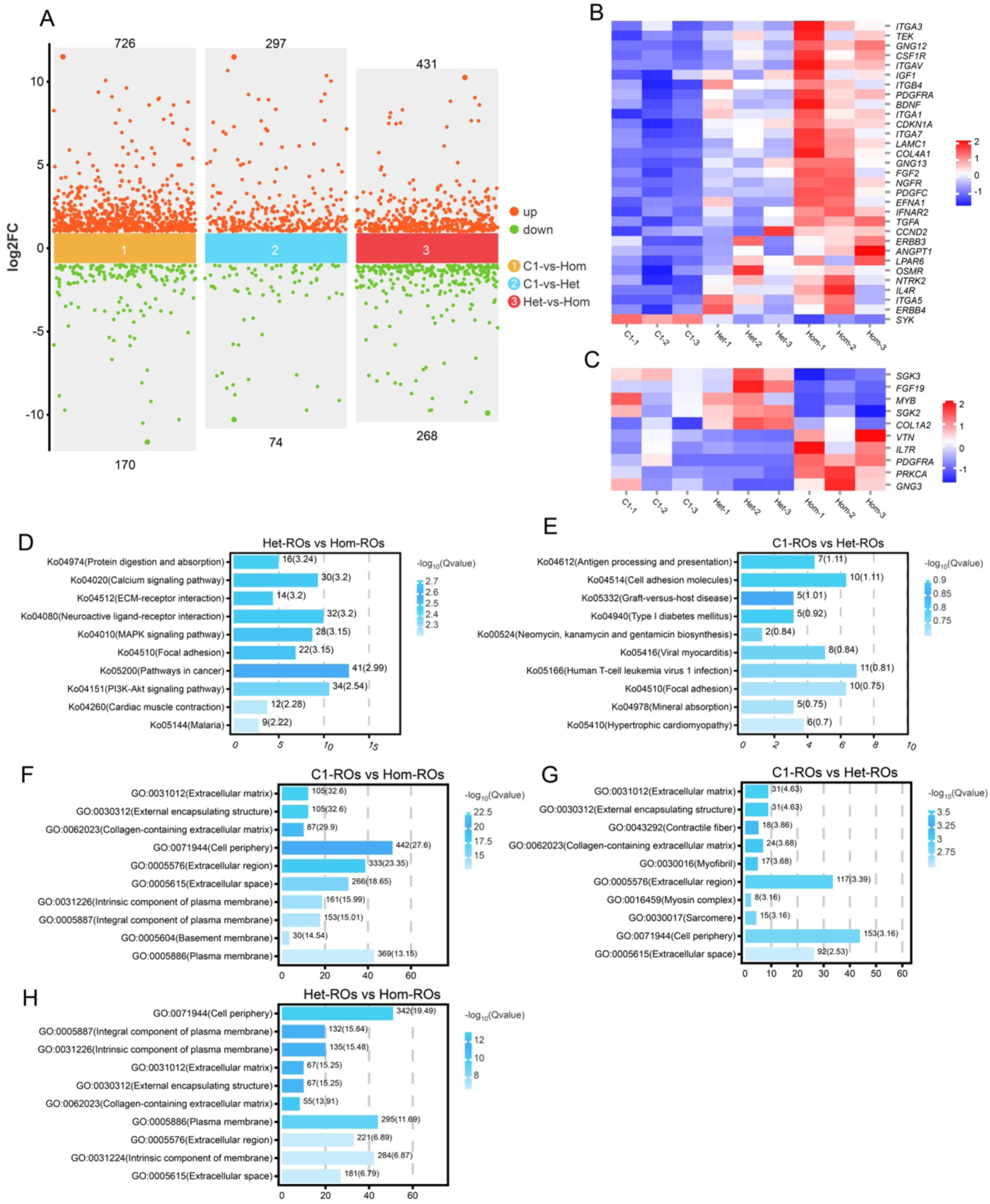
RNA-seq analysis reveals changes induced by the *PROM1* mutation in patient-specific photoreceptors. **(A)** Pairwise comparisons among C1-ROs, Het-ROs, and Hom-ROs reveal differentially expressed genes (DEGs), with counts indicated above and below each line. DEGs were defined by an absolute log₂(fold change) > 1.5 and an adjusted P-value < 0.05. **(B and C)** Supplementary to Figure 3G. Panel (B) shows DEGs in the PI3K-Akt signaling pathway (ko04151) unique to the C1-ROs vs. Hom-ROs comparison, while panel (C) shows those unique to the Het-ROs vs. Hom-ROs comparison. **(D and E)** KEGG pathway analysis of the top 10 significantly altered pathways between Het-ROs vs Hom-ROs (D) and C1-ROs vs Het-ROs (E). **(F-H)** GO enrichment analysis results from pairwise comparisons among C1-ROs, Het-ROs, and Hom-ROs.

**Figure S11.**
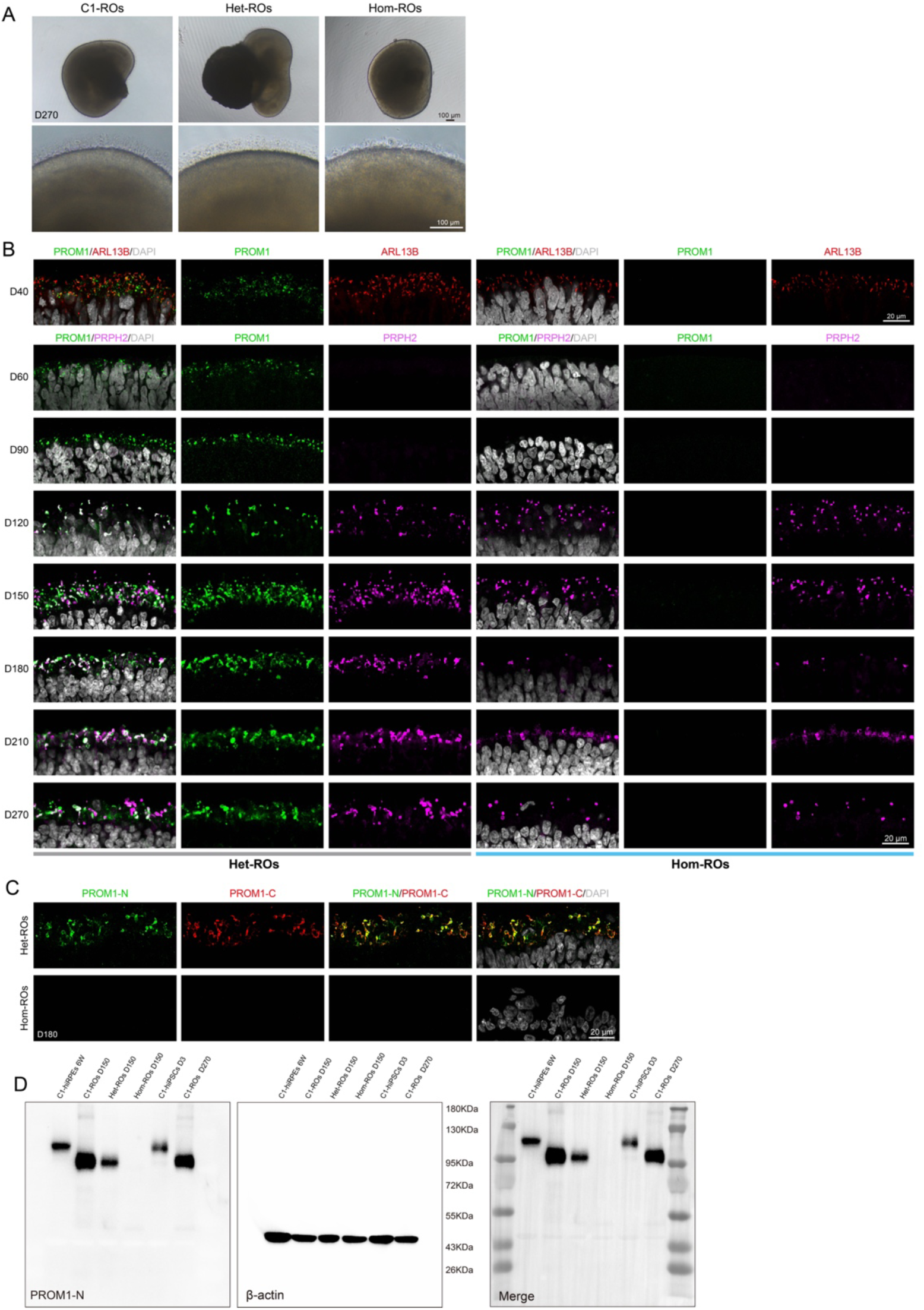
Mutant PROM1 protein is undetectable in patient-specific ROs. **(A)** Bright-field images of ROs at D270. Scale bar, 100 µm. **(B)** Immunostaining shows expression of PROM1, ARL13B, and PRPH2 across differentiation timepoints (D40–D270) in Het-ROs and Hom-ROs. Scale bar, 20 µm. **(C)** PROM1 expression in ROs was assessed using PROM1-N and PROM1-C antibodies. **(D)** Original Western blot images corresponding to Figure 3H.

**Figure S12.**
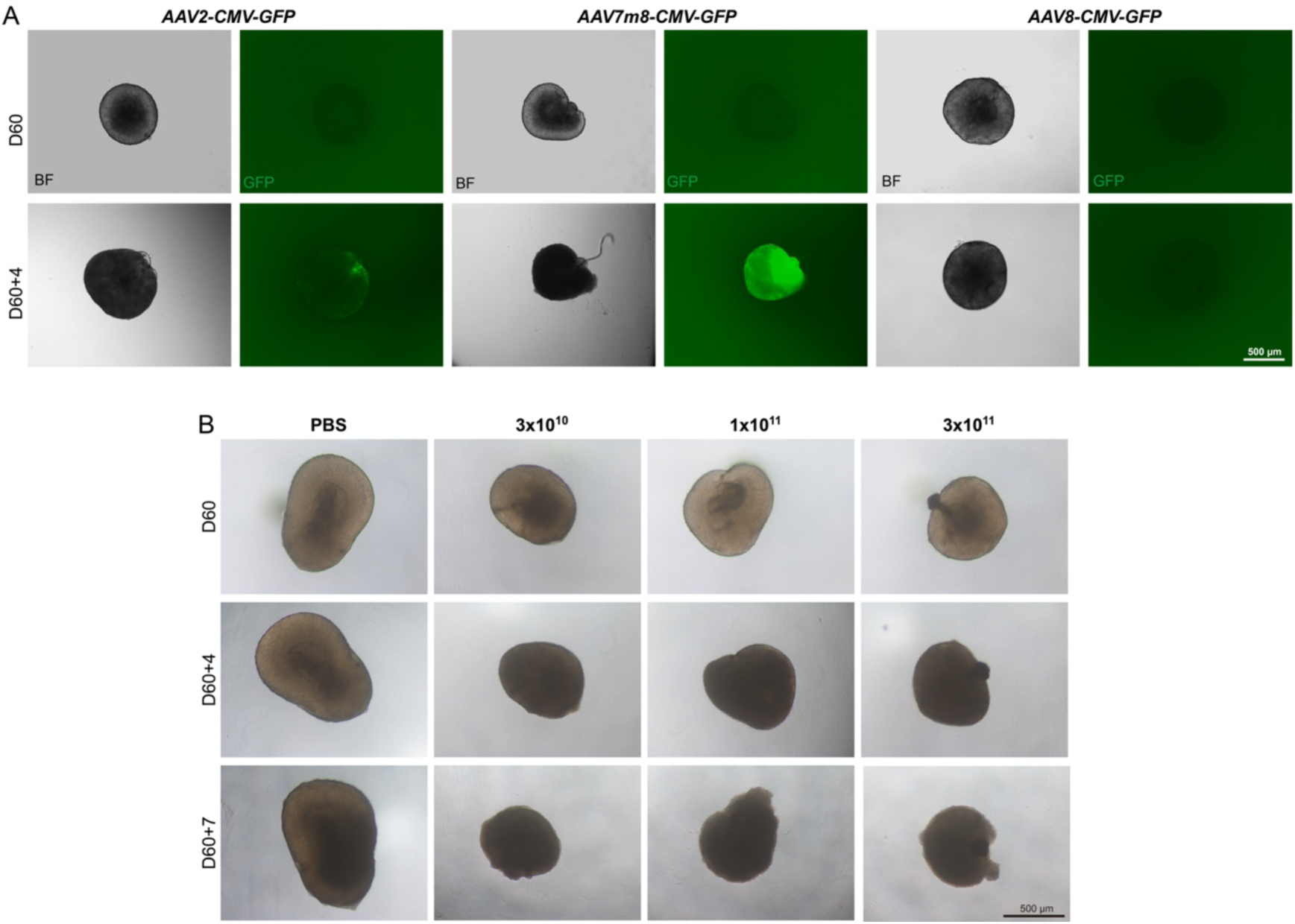
Screening of *AAV* serotypes for efficient transduction in ROs. **(A)** At D60, C1-ROs was infected with *AAV2-CMV-GFP*, *AAV7m8-CMV-GFP*, or *AAV8-CMV-GFP* (1×10¹¹ vg/RO). Changes were assessed by bright-field and GFP fluorescence imaging pre- and post-infection. **(B)** C1-ROs treated with varying concentrations of *AAV7m8-CMV-GFP* (3×10¹⁰, 1×10¹¹, 3×10¹¹ vg/RO) show morphological changes in bright-field images. PBS served as the negative control. Scale bar, 500 µm.

**Figure S13.**
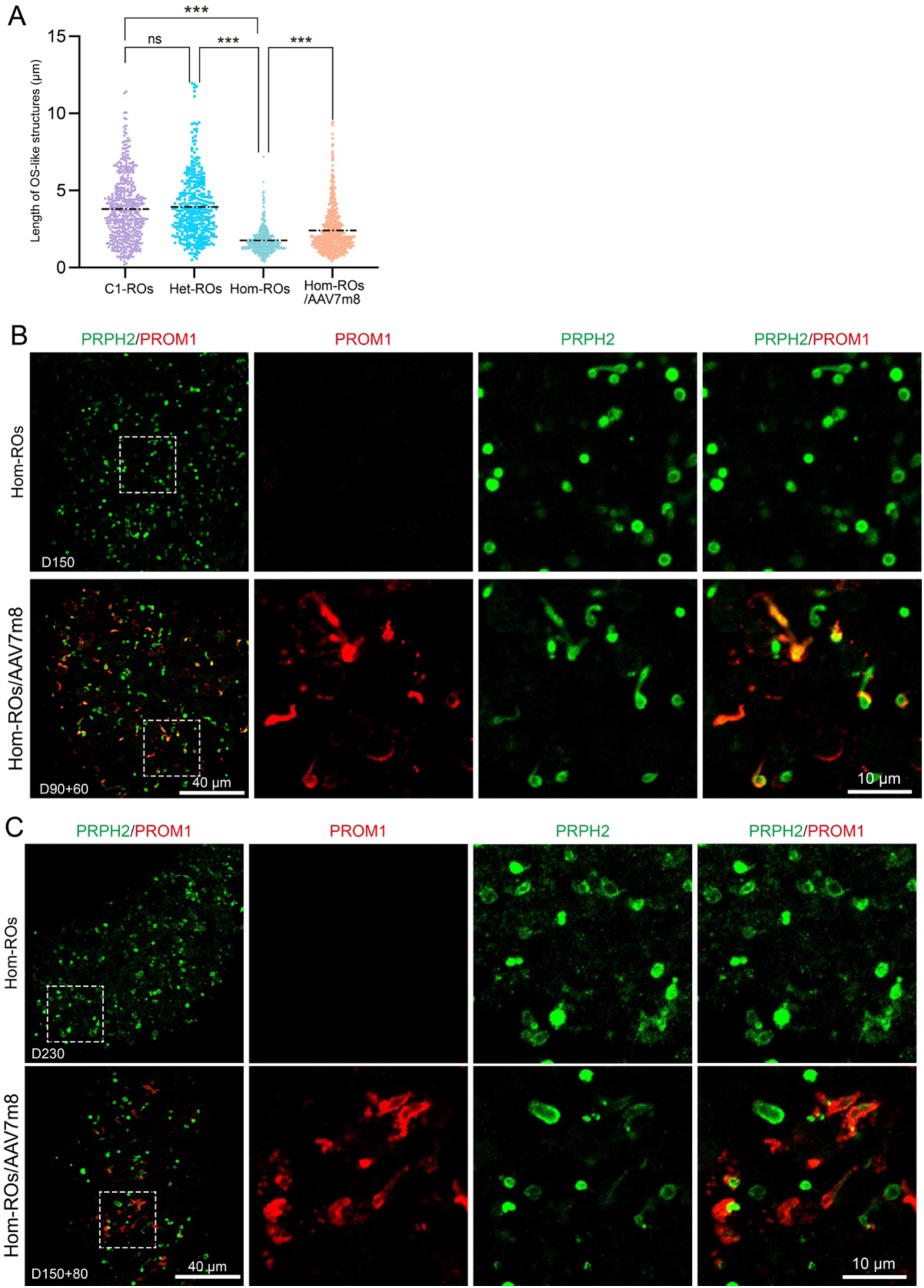
Sustained expression of exogenous PROM1 in OS-like structures. Hom-ROs infected with *AAV7m8-CRXp-hPROM1* was analyzed for PROM1 expression at D90+30 (A), D90+60 (B) and D150+80 (C). Magnified regions indicated by white dashed boxes.

**Figure S14.**
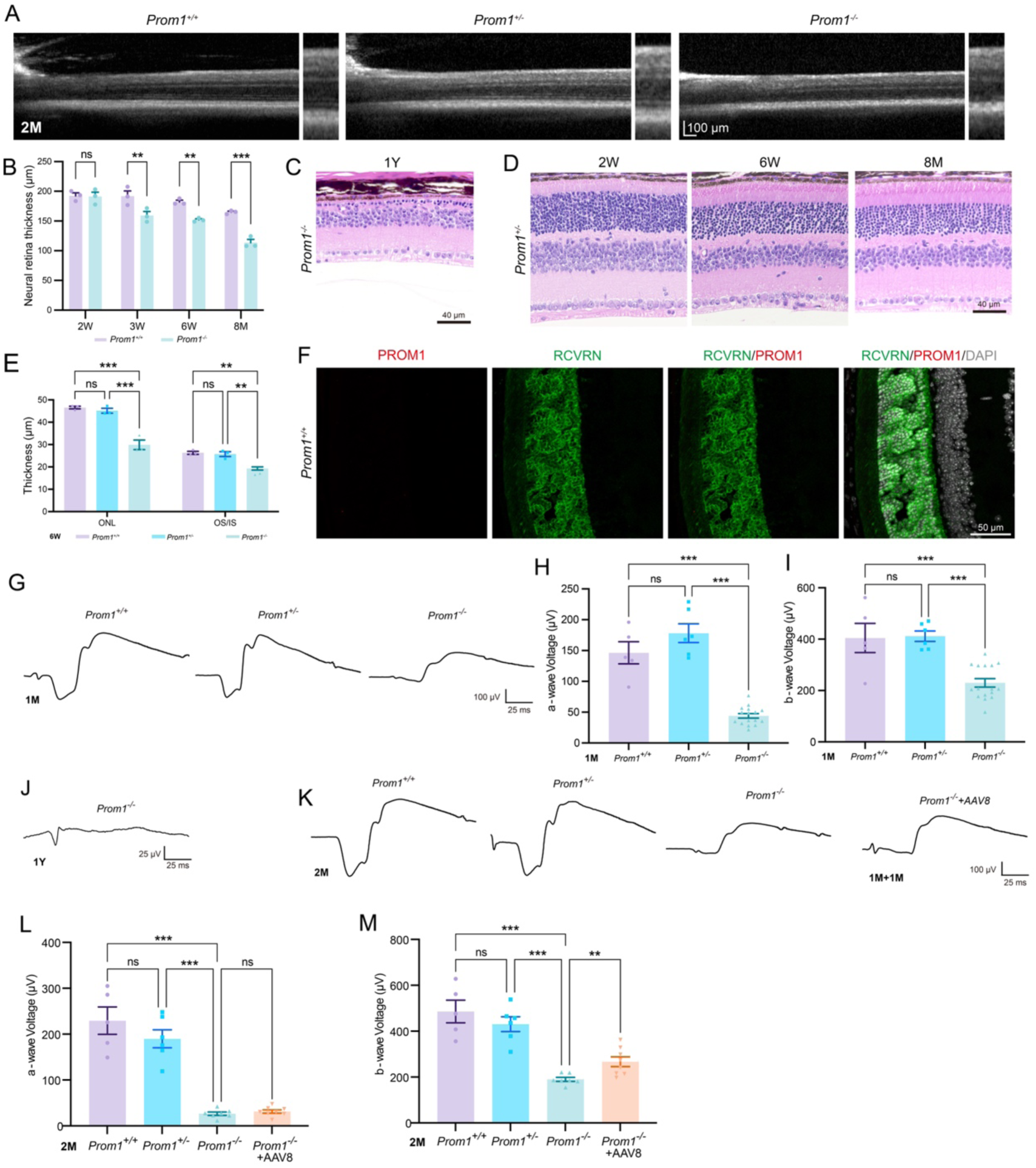
Heterozygous *Prom1* deletion does not lead to retinal degeneration. **(A)** SD-OCT images of 2-month-old mouse retinas. **(B)** Supplementary to Figure 5C. Statistical analysis of retinal thickness changes from 2 weeks to 8 months in *Prom1^+/+^* and *Prom1^-/-^* mice. **(C)** H&E staining reveals retinal degeneration in a 1-year-old *Prom1 ^-/-^* mice. Scale bar, 40 µm. **(D)** Retinal morphology of *Prom1^+/−^* mice at 2 weeks, 6 weeks, and 8 months. Scale bar, 40 µm. **(E)** Comparison of ONL and IS/OS thickness in 6-week-old *Prom1^+/+^*, *Prom1^+/-^*, and *Prom1^-/-^* mouse retinas. **(F)** PROM1-N staining in *Prom1^+/+^* retinas. Scale bar, 50 µm. **(G–I)** Scotopic 3.0 recordings in PM1 mice. Panel (G) displays representative waveforms. The a- and b-wave amplitudes are quantified in (H) and (I). Groups: *Prom1^+/+^* (n=5), *Prom1^+/-^* (n=6), and *Prom1^-/-^* (n=16). **(J)** Extinguished fERG response in 1-year-old *prom1^-/-^* mice. **(K–M)** Representative waveforms in PM2 mice (J). The amplitudes of the a- and b-wave are detailed in (K) and (L). *Prom1^+/+^* (n=5), *Prom1^+/-^* (n=6), *Prom1^-/-^* (n=7), and *Prom1^-/-^+AAV8-hPROM1* (n=8). *Error bars represent mean ± S.E.M.; ns, not significant (P ≥ 0.05); **P < 0.01; ***P < 0.001*.

**Figure S15.**
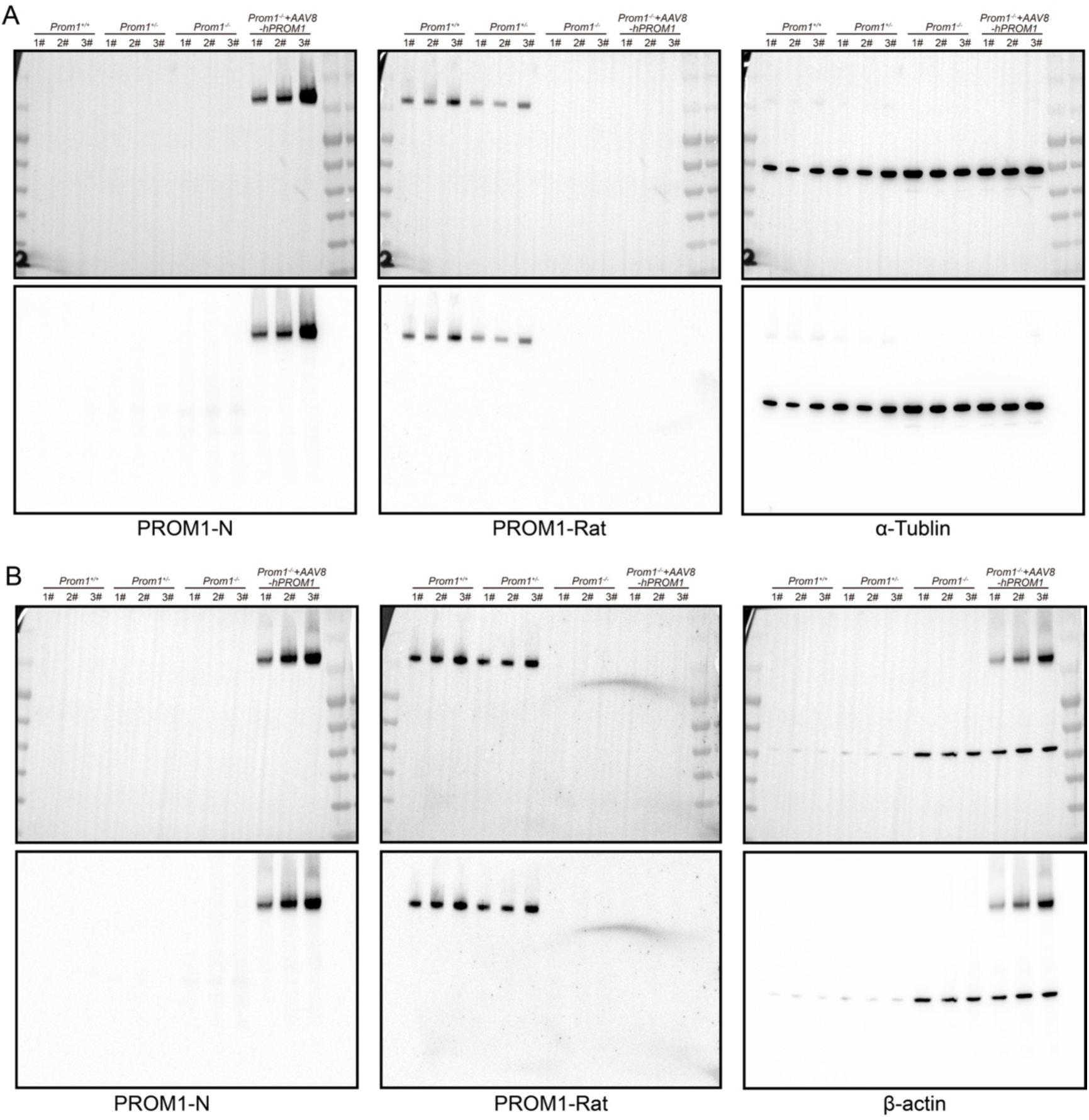
Expression of exogenous human PROM1 in *AAV8-CRXp-hPROM1* infected retina of *Prom1^-/-^* mice. **(A and B)** Western blots of PROM1 in mouse retinas, showing original image (A) and technical replicate (B). β-Actin as loading control.

**Table S1.** Antibodies used in this study.

| Antibody | Vendor | Cat. No. | RRID | For IF | For WB | Host |
| --- | --- | --- | --- | --- | --- | --- |
| **First antibodies** |  |  |  |  |  |  |
| Anti-PROM1 (PROM1-N) | Invitrogen | MA1-219 | AB_2725113 | 1: 200 | 1:1000 | Mouse |
| Anti-PROM1 (PROM1-C) | Abcam | ab19898 | AB_470302 | 1: 100 | 1:1000 | Rabbit |
| Anti-PROM1 (PROM1-Rat) | Invitrogen | 14-1331-95 | AB_2864986 | 1: 200 | 1:1000 | Rat |
| Anti-ARL13B | Proteintech | 17711-1-AP | AB_2060867 | 1: 200 | - | Mouse |
| Anti-PRPH2 | Proteintech | 18109-1-AP | AB_10665364 | 1:200 | - | Rabbit |
| Anti-Arrestin 3 | Novus | NBP1-37003 | AB_2060085 | 1:200 | - | Goat |
| Anti-Rhodopsin | Abcam | ab5417 | AB_304874 | 1:200 | - | Rabbit |
| Anti-L/M-opsin | Kind gift from Dr. Jeremy Nathans |  |  | 1:500 | - | Rabbit |
| Anti-S-opsin | Kind gift from Dr. Jeremy Nathans |  |  | 1:500 | - | Rabbit |
| Anti-MCT1 | Proteintech | 20139-1-AP | AB_2878645 | 1:300 | - | Rabbit |
| Anti-ZO-1 | Invitrogen | 33-9100 | AB_87181 | 1:400 | - | Mouse |
| Anti-VSX2 | Millipore | ab9016 | AB_2216009 | 1:200 | - | Sheep |
| Anti-Recoverin | Millipore | ab5585 | AB_2253622 | 1:500 | - | Rabbit |
| Anti-CRALBP | Abcam | ab15051 | AB_2269474 | 1:500 | - | Rabbit |
| Anti-RBPMS | Novus | GTX118619 | AB_10720427 | 1:500 | - | Rabbit |
| Anti-OTX2 | Abcam | ab21990 | AB_776930 | 1:500 | - | Rabbit |
| Anti-PROX1 | Millipore | AB5475 | AB_177485 | 1:2000 | - | Rabbit |
| Anti-AP2α | DSHB | 3B5 | AB_528084 | 1:35 | - | Rabbit |
| Anti-PKCα | Abcam | ab32376 | AB_777294 | 1:2000 | - | Rabbit |
| Anti-β-actin | Beyotime | AF2811 | AB_3668910 | - | 1:5000 | Mouse |
| Anti-α-Tubulin | Beyotime | AF2827 | AB_3064710 | - | 1:2000 | Mouse |
| **Second antibodies** |  |  |  |  |  |  |
| Donkey Anti-Mouse (555) | Invitrogen | A31570 | AB_2536180 | 1:500 | - | Donkey |
| Donkey Anti-Rabbit (555) | Invitrogen | A31572 | AB_162543 | 1:500 | - | Donkey |
| Donkey Anti-Goat (555) | Invitrogen | A21432 | AB_141788 | 1:500 | - | Donkey |
| Donkey Anti-Mouse (488) | Invitrogen | A21202 | AB_141607 | 1:500 | - | Donkey |
| Donkey Anti-Rabbit (488) | Invitrogen | A21206 | AB_2535792 | 1:500 | - | Donkey |
| Donkey Anti-Mouse (647) | Invitrogen | A31571 | AB_162542 | 1:500 | - | Donkey |
| Donkey Anti-Rabbit (647) | Invitrogen | A31573 | AB_2536183 | 1:500 | - | Donkey |
| HRP-Goat Anti-Mouse IgG | Beyotime | A0216 | AB_2860575 | - | 1:2000 | Goat |
| HRP-Goat Anti-Rabbit IgG | Beyotime | A0208 | AB_2892644 | - | 1:1000 | Goat |
| HRP-Goat Anti-Rat IgG | Beyotime | A0192 | AB_2939016 | - | 1:1000 | Goat |

**Table S2.**
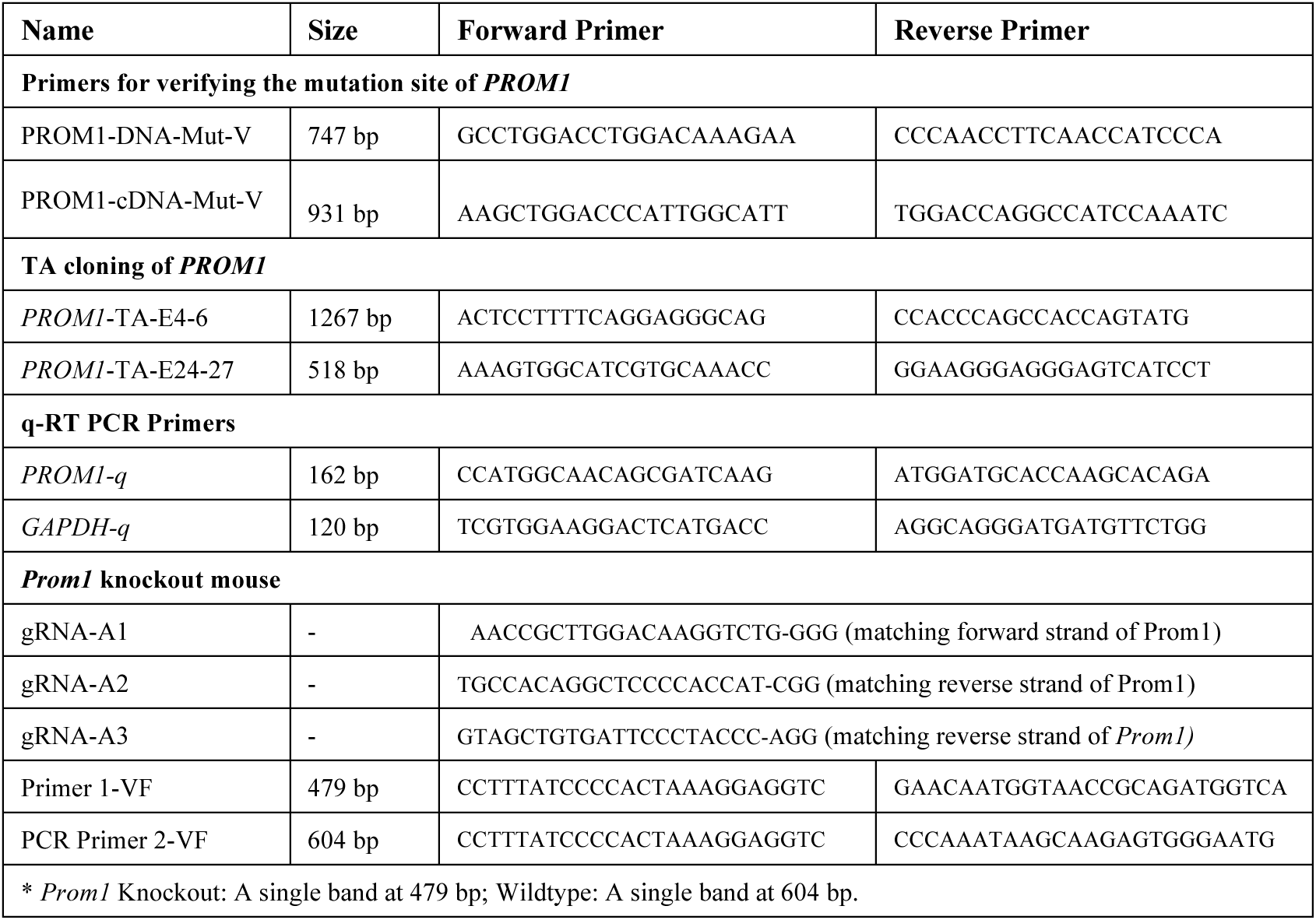
Primer used in this study.

